# Differential tissue deformability underlies shape divergence of the embryonic brain and spinal cord under fluid pressure

**DOI:** 10.1101/2024.01.12.575349

**Authors:** Susannah B.P. McLaren, Shi-Lei Xue, Siyuan Ding, Alex Winkel, Oscar Baldwin, Shreya Dwarakacherla, Kristian Franze, Edouard Hannezo, Fengzhu Xiong

**Affiliations:** Wellcome Trust / CRUK Gurdon Institute, University of Cambridge, Cambridge, UK; Department of Physiology, Development and Neuroscience, University of Cambridge, Cambridge, UK; Institute of Science and Technology Austria, Klosterneuburg, Austria; School of Engineering, Westlake University, Hangzhou, China; Friedrich-Alexander-Universität Erlangen-Nürnberg, Germany; Max-Planck-Zentrum für Physik und Medizin, Germany

## Abstract

An expanded brain enables the complex behaviours of vertebrates that promote their adaptation in diverse ecological niches^1–3^. Initial morphological differences between the brain and spinal cord emerge as the antero-posteriorly patterned neural plate folds to form the neural tube^4–7^ during embryonic development. Following neural tube closure, a dramatic expansion of the brain diverges its shape from the spinal cord^8^, setting their distinct morphologies for further development^9,10^. How the brain and the spinal cord expand differentially remains unclear. Here, using the chicken embryo as a model, we show that the hindbrain expands through dorsal tissue thinning under a positive hydrostatic pressure from the neural tube lumen^11,12^ while the dorsal spinal cord shape resists the same pressure. Using magnetic droplets and atomic force microscopy, we reveal that the dorsal tissue in the hindbrain is more fluid than in the spinal cord. The dorsal hindbrain harbours more migratory neural crest cells^13^ and exhibits reduced apical actin and a disorganised laminin matrix compared to the dorsal spinal cord. Blocking the activity of neural crest-associated matrix metalloproteinases inhibited dorsal tissue thinning, leading to abnormal brain morphology. Transplanting early dorsal hindbrain cells to the spinal cord was sufficient to create a region with expanded brain-like morphology including a thinned-out roof. Our findings open new questions in vertebrate head evolution and neural tube defects, and suggest a general role of mechanical pre-pattern in creating shape differences in epithelial tubes.

## MAIN

The neural tube gives rise to the brain and spinal cord during vertebrate embryonic development. In avian embryos, the volumetric expansion of the brain over a relatively stable spinal cord takes place shortly after neural tube closure (Figure 1A) and depends on the hydrostatic pressure from the neural tube lumen^14,11^. In the stages prior to brain expansion (before Hamburg-Hamilton (HH) stage 13^8^), the neural tube has large openings at both the anterior and posterior ends that allow fluid to move between the tube lumen and its surroundings (Figure 1B). These openings narrow and close concomitant with the onset of brain expansion around HH13^15^, such that the neural tube lumen becomes a closed and continuous fluid filled cavity (Figure S1A). Consistent with pioneering studies^16^, we recorded a positive internal lumen pressure of ∼15Pa in the initial stages of brain expansion (HH13-15), which increased to ∼25Pa as brain expansion progressed (HH19-21. Figure 1C and S1B). To assess how neural tube morphology changes as internal lumen pressure increases, we measured tissue thickness around the circumference of the lumen in cross sections of the hindbrain and spinal cord prior to and after brain expansion (HH11 and HH16, respectively. Figure 1D-F). Dorsal hindbrain tissue approximately halved in thickness and formed a single-cell thick epithelium concomitant with brain expansion, whilst little dorsal thinning or tissue shape change was observed in the spinal cord (Figure S1C and D).

**Figure 1.**
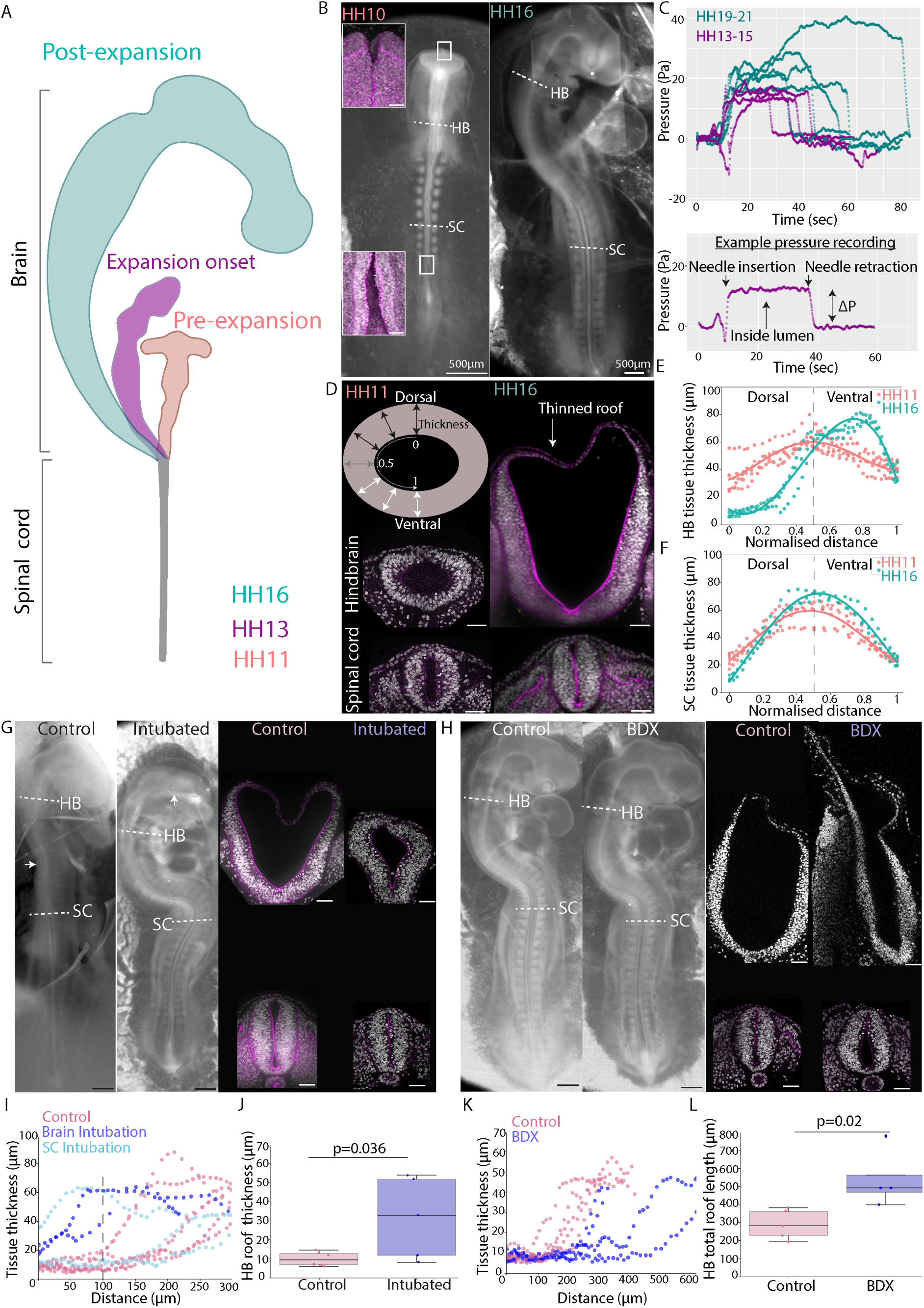
Neural tube lumen pressure drives hindbrain and spinal cord shape divergence. (A) Schematic depicting brain expansion during chicken embryo development, Hamburger and Hamilton stages of development are given. (B-F) Lumen pressure increase and dorsal tissue thinning occur concomitantly with brain expansion. (B) Bright-field images of chicken embryos prior to brain expansion (HH10) and following brain expansion (HH16) and confocal microscopy images of the early anterior and posterior neuropore at the level of the white boxes. Actin is shown in magenta throughout. (C) Pressure readings obtained by inserting a microneedle connected to a pressure sensor into the neural tube lumen in embryos at the onset of (n=4) and following brain expansion (n=5). (D) Schematic showing tissue thickness measurement approach and confocal microscopy images of neural tube cross-sections taken at the level of the dashed white lines in (B). In some cases the thin dorsal tissue collapsed downwards following fixation. (E) Apical-basal tissue thickness along the hindbrain lumen circumference, n=6 HH11 embryos, n=3 HH16 embryos. (F) Apical-basal tissue thickness along the spinal cord lumen circumference, n=6 HH11 embryos, n=3 HH16 embryos. (G-L) Pressure decrease and increase experiments. (G) Bright-field images of control and intubated embryos at ∼20 hours post-intubation and confocal images of corresponding hindbrain and spinal cord cross-sections. (H) Bright-field images of control and β-D-xyloside (BDX) treated embryos ∼20h post treatment and corresponding confocal images of hindbrain and spinal cord cross-sections. (I) Tissue thickness along the hindbrain circumference moving from dorsal to ventral in control, anterior intubated and spinal cord intubated embryos. (J) Hindbrain roof thickness (averaged over the first 100µm from the dorsal midpoint in control (n=6), anterior intubated (n=2) and spinal cord intubated (n=3) embryos. (K) Tissue thickness along the hindbrain circumference in control and BDX treated embryos. (L) Hindbrain total roof length in control (n=5) and BDX treated (n=4) embryos. Black scale bars are 500µm, white scale bars are 50µm unless otherwise stated.

To test the role of lumen pressure in dorsal thinning, we intubated^14^ embryos in the brain or spinal cord just prior to brain expansion (HH11), leaving the tube lumen connected to the embryo’s surroundings to equate the pressure (Figure S1E). Although we observed an increase in hindbrain cross-sectional area, in line with proliferation and tissue growth, dorsal hindbrain thinning and expansion were inhibited in intubated embryos following 20 hours of incubation (Figure 1G, I and J). Not only brain-intubated but also a spinal cord-intubated embryo showed inhibited hindbrain thinning, confirming that the lumen is connected along the anterior-posterior axis of the neural tube and the tissues experience the same hydrostatic pressure. Conversely, treating embryos with β-D-xyloside (BDX) ^17^, which has been found to osmotically increase lumen pressure by ∼30% ^12^, led to an increase in expansion and extension of the thinned dorsal roof in the hindbrain and a moderately enlarged spinal lumen with no dorsal thinning (Figure 1H, K and L). Together these findings demonstrate that neural tube lumen pressure drives dorsal tissue thinning in the hindbrain but is resisted in the dorsal spinal cord, leading to brain expansion relative to the spinal cord.

The differential thinning of dorsal tissue between the hindbrain and the spinal cord may result from a few scenarios. First, different initial tissue curvature at the onset of pressure increase (Figure 1D) can lead to diverging dynamics of thinning and volume expansion. Generically, forces exerted on a tubular tissue are proportional to both the luminal fluid pressure and the tissue radius. Driven by the same pressure, the tubular region with a greater initial radius (lower curvature) would then display larger tissue forces and expansion (Figure 2A), which could be consistent with the shape dynamics observed in the hindbrain and spinal cord. To experimentally test this possibility, we modified spinal cord lumen shape prior to brain expansion by performing *in ovo* surgeries to remove somites and anterior presomitic mesoderm on either side of the spinal cord (Figure 2B). This operation releases tissue confinement from not only the somites (Figure S2A) but also the dorsal non-neural ectoderm as it was cut through during somite ablation. After ∼20 hours of incubation, these embryos healed and exhibited reduced spinal cord dorsal curvature and in one case clear rounding of the spinal cord cross section (Figure 2C, S2B-C). However, this did not lead to the formation of a thinned-out roof, as was observed in the hindbrain (Figure S2D). These results suggest that the higher curvature of the spinal cord dorsal region is not sufficient to explain the lack of dorsal tissue thinning in the spinal cord. At the same time, this experiment ruled out a second alternative model in which local confinement from neighbouring tissues (such as somites and the presomitic mesoderm) might modulate neural tube shape (^18^ and Figure S2A) by constraining lumen expansion of the spinal cord relative to the hindbrain. This led us to explore a third hypothesis: that the differential expansion of the brain and spinal cord was due to the dorsal tissue of the hindbrain being more deformable than that of the spinal cord, enabling local thinning despite a spatially uniform hydrostatic pressure. To test whether the material properties of the hindbrain and spinal cord dorsal tissue differed before the onset of increased lumen pressure, we injected ferrofluid oil droplets into the forebrain of HH11 embryos and positioned the droplets in either the hindbrain or spinal cord using a magnetic field (Figure 2D). We then observed the resulting tissue deformation from the pressure generated by the droplets as they try to round up due to their surface tension after removal of the magnetic field (Figure 2E). Droplets were injected either at equal volumes (2nl) or at a ratio to account for the difference in initial lumen size (4nl in the larger hindbrain lumen, 1nl in the smaller spinal cord lumen). In both cases the droplets showed a higher curvature at the droplet-lumen interface in the spinal cord, indicating a higher pressure was exerted on the spinal cord tissue than the hindbrain by this mechanical perturbation^19^ (Figure S2E). Still, the thickness change of dorsal tissue at the droplet site was greater in the hindbrain than in the spinal cord (Figure 2F), indicating that hindbrain dorsal tissue was more deformable. Furthermore, observing droplet shape change over time revealed a slow rounding process occurring over hours (Movies S1 and S2), orders of magnitude longer than would be expected from droplet surface tension and viscosity alone, which provided an opportunity to estimate the effective long-term viscous response of the surrounding tissue (Figure S2F). To do this, we modelled the neural tube as a viscoelastic Maxwell medium with differing geometry between the brain and spinal cord (supplemental text). By fitting droplet shape dynamics to the model, we found that the estimated hindbrain viscosity was much lower than the spinal cord and that a lumen pressure of 10’s of Pa is capable of driving hindbrain expansion over long timescales, consistent with our pressure measurements (Figure 1C). As an independent validation of our findings on tissue mechanical properties, we used an atomic force microscopy (AFM) cantilever to indent the dorsal region of the hindbrain and spinal cord (Figure 2G) in the same embryos and measured tissue relaxation in response to a sustained force (50 or 75nN). We found that the dorsal hindbrain had a higher fluidity than that of the spinal cord prior to brain expansion on short timescales, whilst no significant difference in instantaneous elastic modulus was detected (Figure 2H and I). While *in vivo* AFM measurements may contain a contribution of differences from other non-local structures such as the extracellular matrix (ECM) anchoring the neural tube, these results are consistent with our droplet experiments. Furthermore, incorporating a less viscous dorsal region into our neural tube model generated tissue thinning and expansion dynamics consistent with experimental observations of hindbrain morphogenesis (supplemental text, section 4). Together, these findings support the hypothesis that the dorsal hindbrain deforms more than the dorsal spinal cord under lumen pressure through viscous thinning over time. To investigate the mechanism of dorsal tissue deformability, we explored the cellular organisation of the tissue at stages leading up to brain expansion (HH9-12). The dorsal neural tube tissue is occupied by neural crest cells that reorganise their actomyosin cytoskeleton as they undergo an epithelial-to-mesenchymal transition (EMT) and delaminate from the tube dorsal surface as the walls of the neural plate fuse dorsally ^20,21,22^. The dynamics of neural crest delamination between cranial and trunk regions is known to differ ^23,24^, making them an exciting candidate for underlying tissue intrinsic differences between the dorsal hindbrain and spinal cord. We observed clear apical organisation of actin and myosin within the dorsal spinal cord in HH12 embryos, however this organisation was largely absent in the neural crest domain of the hindbrain (Figure 3A-C). This change in cytoskeleton organisation indicated that a transition away from an epithelial cell state to a more mesenchymal cell state was occurring in the dorsal hindbrain. A key player in this process is the zinc-finger transcription factor Snail2 which is expressed in the pre-migratory and early migrating neural crest ^25,26^. Using Snail2 as a marker, we observed that the dynamics of neural crest delamination and migration at the dorsal surface of the neural tube differed between the hindbrain and the spinal cord leading up to brain expansion (HH9-12), with Snail2+ cells migrating in a cell-by-cell fashion in spinal cord regions, whilst those in the hindbrain migrate in a collective wave (Figure 3D and S3A). Neural crest cells utilise matrix metalloproteases (MMPs) to remodel the surrounding ECM ^27,28^. ECM architecture plays an important role in determining tissue material properties, with higher levels of ECM organisation conferring greater stiffness (or elasticity) and more disordered matrix organisation leading to softening (or a more viscous behaviour) ^29,30^, which contributes to long-term deformability.

**Figure 2.**
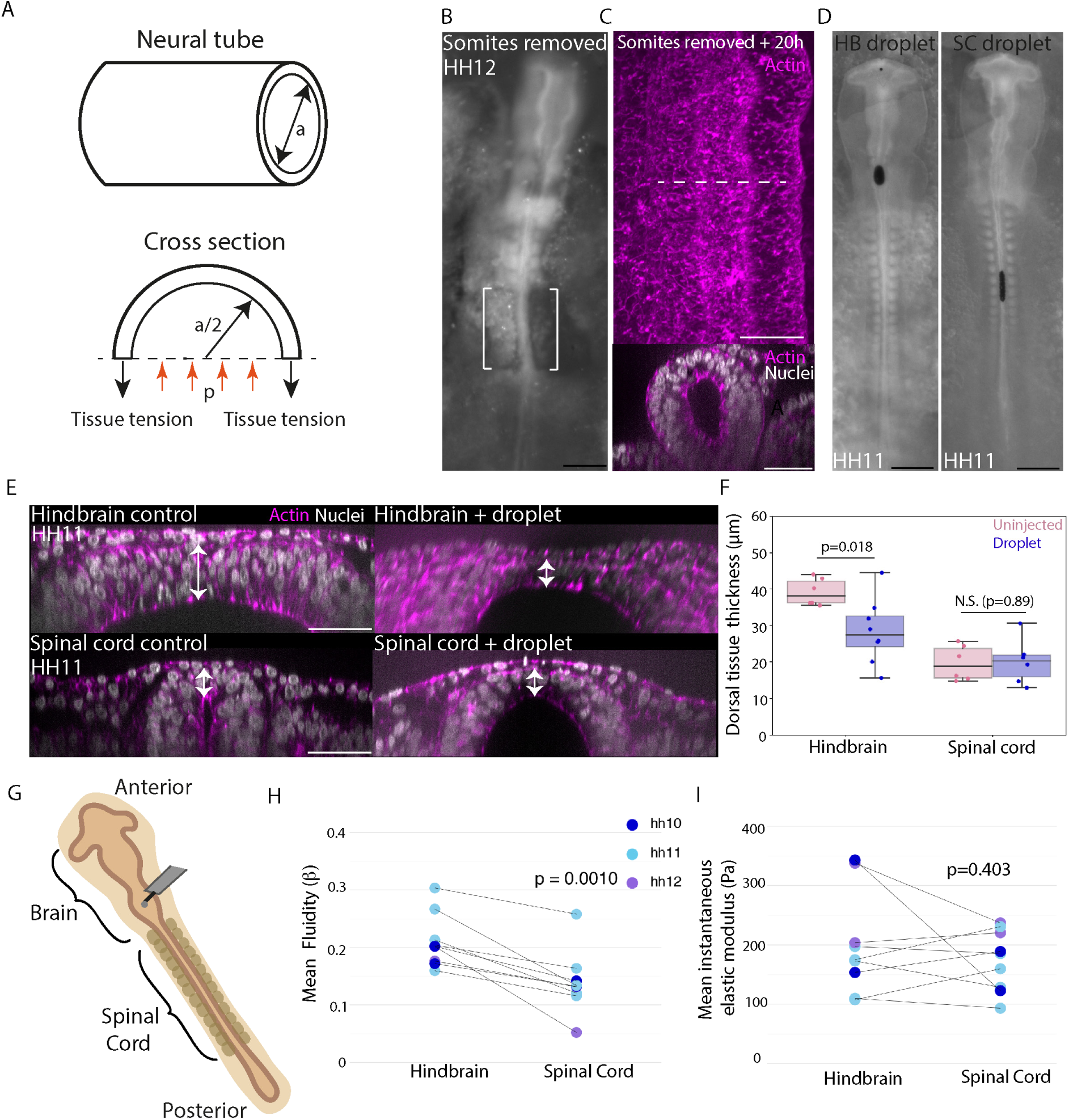
The dorsal hindbrain is more deformable than the dorsal spinal cord prior to brain expansion. (A) Schematic illustrating that forces exerted on a tubular tissue are generically proportional to lumen pressure and tissue radius. (B-C) Lateral tissue removal experiments. (B) Bright-field image of a pre-brain expansion stage embryo with posterior somites and anterior presomitic mesoderm removed (n=3, control n=3). (C) Confocal images of the dorsal surface view and cross-section view of the spinal cord in a ∼20h post somite-removal embryo. (D-F) Dorsal tissue deformation under ferrofluid droplet injection. (D) Widefield images of HH11 stage embryos with ferrofluid droplet injections in either the hindbrain or spinal cord lumen. (E) Confocal images showing cross-sectional views of the spinal cord and hindbrain in HH10-11 stage uninjected embryos and embryos with ferrofluid droplets injected into either the hindbrain or spinal cord. White arrows indicate thickness measurements (acquired at the thinnest dorsal point observable). (F) Quantification of hindbrain and spinal cord roof thickness in uninjected (n=6 hindbrain, n=6 spinal cord) and droplet injected (n=8 hindbrain, n=6 spinal cord) embryos. (G-I) Atomic force microscopy experiments. (G) Schematic depicting atomic force microscopy measurement set up. (H) Mean Beta value (corresponding to fluidity) of the hindbrain and spinal cord in HH10-12 embryos (n=9). (I) Mean instantaneous elastic modulus of the hindbrain and spinal cord in HH10-12 embryos (n=9). Black scale bars are 500µm. White scale bars are 50µm.

**Figure 3.**
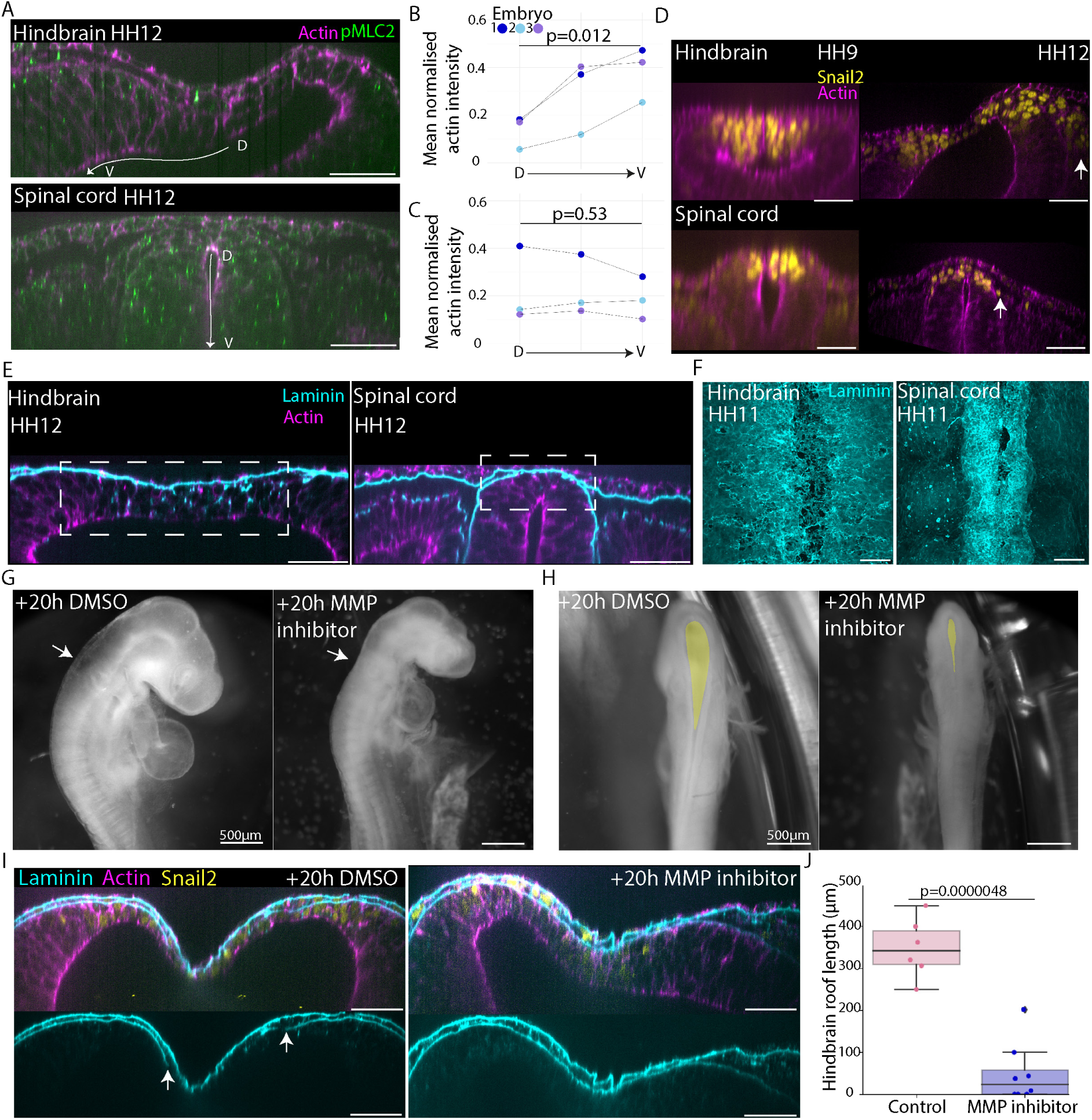
Differential neural crest cell behaviour underlies the acquisition of a thinned out and expanded dorsal hindbrain roof. (A-C) Actomyosin localisation in the dorsal hindbrain and spinal cord. (A) Confocal images of actin and phosphorylated myosin light chain kinase II in the hindbrain and spinal cord of HH12 stage embryos. (B) Quantification of apical actin signal intensity along the lumen circumference in the hindbrain of HH11 and HH12 stage embryos. (C) Quantification of apical actin signal intensity along the lumen circumference in the spinal cord of HH11 and HH12 stage embryos. (D) Premigratory neural crest in the hindbrain and spinal cord. Confocal images showing Snail2+ cell migration in the hindbrain and spinal cord cross sections of embryos progressing towards brain-expansion stages (HH9 to HH12). (E-F) ECM organisation in the dorsal hindbrain and spinal cord. (E) Confocal images showing laminin and actin organisation in cross sections of the spinal cord and hindbrain of HH12 stage embryos. (F) Confocal images showing laminin organisation at the dorsal surface in a 3D rendering of the spinal cord and hindbrain of HH11 stage embryos. (G-H) ECM remodelling perturbation experiments. (G) Brightfield images of embryos treated with DMSO or MMP inhibitor at HH11 and incubated for 20 hours. White arrows indicate hindbrain at level of otic vesicle. (H) Dorsal views of DMSO and MMP inhibitor treated embryos. Yellow highlight fills the brain dorsal surface. (I) Confocal images showing the dorsal hindbrain of control and MMP inhibitor treated embryos, ∼20 hours post treatment. Bottom panels show laminin surface with white arrows indicating breaks in laminin continuity. (J) Hindbrain roof length obtained by measuring the length of the single-cell thick roof in control (n=6) and inhibitor treated (n=8) embryos. White scale bars are 50µm unless otherwise stated.

This led us to investigate ECM organisation in the hindbrain and the spinal cord as a potential mediator of tissue deformability. Cross-sectional views showed that laminin was redistributed throughout the dorsal hindbrain in the premigratory neural crest domain, whilst comparatively little redistribution was observed in the dorsal spinal cord (Figure 3E). This was associated with a more sparce laminin coating of the hindbrain compared to the spinal cord when viewed from the dorsal side (Figure 3F and S3B and C). These findings suggest that ECM is being actively reorganised and redistributed out of the plane of the basement membrane to a greater extent in the hindbrain.

To test whether ECM remodelling plays a role in dorsal hindbrain thinning, we inhibited MMP activity in embryos prior to brain expansion stages using an MMP-specific small molecule inhibitor (GM6001, 1000µM). Brain expansion was reduced in inhibitor treated embryos and a smaller surface area of the dorsal hindbrain was apparent (Figure 3G-H). In embryos with obvious observable developmental defects (0/6 control, 4/8 inhibitor treated), the hindbrains did not undergo dorsal thinning and in some cases aggregated a mass of Snail2+ cells within the lumen (Figure S3D). In embryos that developed normally overall (6/6 control, 4/8 inhibitor treated), the hindbrains of the treated embryos showed thicker or shorter dorsal roofs than those of the controls (Figure 3I, J and S3D). The laminin surface covering the basal extent of the dorsal hindbrain appeared less disrupted in embryos with inhibited roof thinning (Figure 3I) and the dorsal tissue was still populated with Snail2+ cells, suggesting that neural crest specification continued in the presence of the inhibitor. Together, these data are consistent with the idea that neural crest mesenchymal behaviour and subsequent ECM remodelling is required for thinning and expansion of the dorsal hindbrain. To definitively test the intrinsic ability of early hindbrain dorsal cells to generate a thinned roof under lumen pressure, we performed an *in ovo* graft experiment, moving a small piece of dorsal hindbrain (or spinal cord as a control) tissue from a GFP+ embryo to the pre-cut dorsal region of the spinal cord of a wildtype host before the onset of lumen pressure (Figure 4A). Re-integration of the graft to reform a closed neural tube that would continue normal morphogenesis was expected to be rare given the difficulty of positioning the graft in such a small incision site and maintaining contact between the small graft and the host tissue. After overnight culture, we found 5 embryos with different degrees of hindbrain dorsal tissue integration into the spinal cord, and 2 spinal cord-to-spinal cord controls (Figure 4B, S4A and S4B). The control embryos exhibited a thick dorsal region at the grafted site (Figure 4C, D and S4B, 2/2 embryos). Interestingly, 2 hindbrain-to-spinal cord grafted embryos exhibited dorsal tissue thinning in the grafted region (Figure 4C, D and S4B). In the cleanest graft where the donor tissue completely replaced the host dorsal region, we found a distinctively thinned out dorsal roof and hindbrain-like morphology specific to the grafted region (Figure 4C), with a thicker spinal cord-like host dorsal region present on the anterior and posterior sides of the graft. Cells in the grafted region were Snail2+ (Figure 4E) and the thinned-out roof coincided with loss of apical actin localisation (Figure S4C), suggesting that the grafted region contained premigratory neural crest cells undergoing EMT. This striking ability of Snail2+ dorsal hindbrain cells to form a thinned-out and elongated roof in the spinal cord region suggests that hindbrain premigratory neural crest cells are sufficient to drive dorsal tissue thinning in response to lumen pressure.

**Figure 4.**
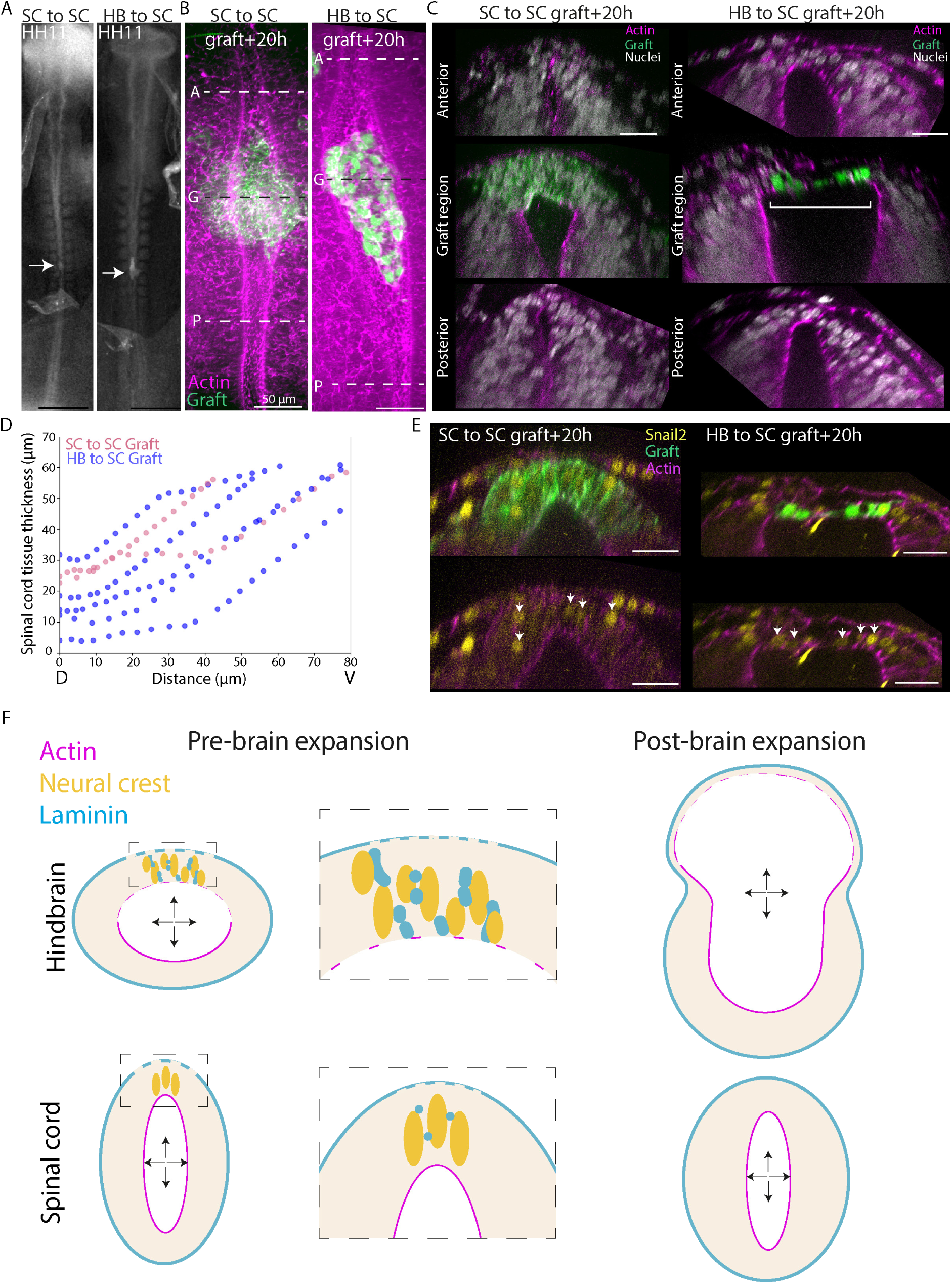
Premigratory neural crest cells in the dorsal hindbrain are sufficient to generate a thinned dorsal roof and neural tube expansion. (A-E) Grafting experiments. (A) Bright-field image of pre-brain expansion stage (HH11) embryos with a spinal cord-to-spinal cord graft or a hindbrain-to-spinal cord graft. (B) Confocal images showing dorsal surface view of spinal cord-to-spinal cord and hindbrain-to-spinal cord graft integration ∼20 hours post-grafting. (C) Confocal images showing cross-sectional views of spinal cord-to-spinal cord and hindbrain-to-spinal cord graft integration ∼20 hours post-grafting. Cross section levels correspond to dashed lines in (B). White bracket highlights the thinned-out roof in the graft region. D) Spinal cord tissue thickness in the region with spinal cord (n=2) or hindbrain (n=5) grafted cells (thickness measurements taken along lumen circumference going from dorsal midpoint and moving ventrally). (E) Confocal images showing Snail2+ cells in graft regions. White arrows indicate Snail2+ cells within the GFP+ graft. (F) Schematic depicting proposed model of brain expansion relative to the spinal cord. Premigratory neural crest cells in pre-brain expansion stage embryos show differences in their behaviour between the hindbrain and spinal cord regions of the neural tube. A greater extent of mesenchymal behaviour and corresponding extracellular matrix remodelling underlies a more deformable dorsal roof in the hindbrain compared to the spinal cord. This allows the hindbrain roof to deform more under internal lumen pressure, driving hindbrain expansion relative to the spinal cord during early embryo development. Black scale bars are 500µm. White scale bars are 25µm unless otherwise stated.

Taken together, our findings support a mechanism where deformability of the neural tube dorsal tissue and ECM organisation underlie the distinct shape changes of the brain and the spinal cord under a shared lumen pressure (Figure 4F). These differences likely result from different behaviours of the neural crest cells, which are known to exhibit different signalling, metabolism and fates between cranial and trunk levels. Our study thus implies a closely coordinated initial developmental step coupling brain expansion and neural crest EMT, both essential contributors to the formation and evolution of the vertebrate head ^31^. We note that these findings are in agreement with classic chimera grafting experiments in which reduced dorsal roof thinning and elongation can be observed in hindbrain regions lacking neural crest cells ^32^. Expanded early brain morphology is highly conserved between chicken and human embryos^33^, and irregularly shaped mesenchymal cells in the dorsal hindbrain have also been observed to contribute to the squamous roof plate in zebrafish embryos ^34^, suggesting that neural crest mediated dorsal tissue fluidisation may contribute to brain expansion in other species. Future work will resolve the detailed molecular and cellular regulation of dorsal neural tissue mechanics and identify genetic and environmental factors that may cause developmental defects associated with aberrant brain expansion through this mechanism. Given the role of lumen pressure in driving morphogenesis of a variety of epithelial tissues and organs ^35,36,37,38^ our findings here in the neural tube imply a general strategy for creating diverse biological shapes via a mechanical property pre-pattern before the onset of changes in fluid pressure.

## ACKNOWLEDGEMENTS

We thank A. Dimitracopoulos, K. Kawaguchi, J. Vidigueira, B. Baum, I. McLaren, D. St Johnston, and members of the Buckley, Scarpa, Steventon, Kawaguchi and Xiong labs for technical assistance and constructive feedback. We thank Ryan Greenhalgh for methods developed to obtain fluidity values from AFM data. We thank Nicola Lawrence, Alex Sossick and Sargon Gross-Thebing from the Gurdon Institute Imaging Facility for microscopy support. This work was supported by a Wellcome Trust / Royal Society Sir Henry Dale Fellowship (215439/Z/19/Z) and an UKRI-EPSRC Frontier Research Grant (EP/X023761/1, originally selected as an ERC Starting Grant) to F.X.

## AUTHOR CONTRIBUTION

S.B.P.M and F.X. conceived the project, designed experiments and wrote the manuscript. S.B.P.M performed all experiments and analysed the data. S.X and E.H. developed the theoretical model and wrote the supplementary theory note. S.D, O.B, S.D and F.X contributed to embryo experiments. K.F. and A.W. designed and performed AFM experiments.

## MATERIALS & METHODS

### Egg lines and embryo culture

Wild type fertilised chicken eggs were obtained from MedEgg Inc., GFP+ eggs (Cytoplasmic ^39^and Membrane^40^) were obtained from the National Avian Research Facility (NARF) at University of Edinburgh. Eggs were kept in a fridge at 14°C for storage and in a 37.5°C humidified (40%-60%) incubator (Brinsea) during incubation. No animal protocol under the UK Animals (Scientific Procedures) Act 1986 was required for the chicken embryo stages investigated (incubated under 2 weeks, or 2/3 of the gestation time). *Ex ovo* culture was performed using a modified EC culture^41^ protocol as described^42^. For *in ovo* operations, egg shells were windowed with surgical scissors after removing some albumen. A small volume of Chinese calligraphy ink (YiDeGe, Black) was then injected underneath the embryo to help with visualisation. After performing surgical, mechanical or chemical perturbations, eggs were resealed with clear plastic tape and returned to incubation.

### Pressure measurement

Pressure measurements were performed as detailed in ^43^. Briefly, a glass microneedle connected to a pressure sensor was inserted into the neural tube lumen of chicken embryos at varying stages of development. Pressure readings were continuously recorded to include the pressure just outside the lumen, after insertion into the lumen, and after retraction from the lumen for each experiment. The pressure difference between the outside and inside of the lumen was taken as the neural tube lumen pressure. Measurements where no stable pressure reading could be obtained inside the neural tube lumen, likely due to needle clogging by tissue debris during the insertion, were excluded from analysis.

### Pressure modulation

Intubation experiments were performed by inserting an open-ended short glass capillary tube into the forebrain or spinal cord region of HH10-12 stage embryos. Control experiments were performed by either inserting a glass capillary into tissues just neighbouring the neural tube or inserting a very narrow glass capillary and not submerging it in fluid with the embryo. β-D-xyloside (BDX) experiments were performed by adding either DMSO or BDX to embryo culture medium at a final concentration of 2mM in L15 media. In all cases embryos were cultured overnight *ex ovo*. Only embryos that displayed continued development were included in the analysis.

### Assessment of lumen continuity

Fluorescently labelled Dextran was injected into the neural tube lumen of HH11-13 stage embryos. The fluorescent signal was imaged following injection to assess how far the dye travelled along the neural tube lumen. The dye was observed to spread anteriorly into the brain region and posteriorly into the posterior spinal cord when injected into the spinal cord lumen mid-way along the trunk.

### Removal of confining tissues

Posterior somites and anterior presomitic mesoderm were removed from embryos *in ovo* using a sharpened tungsten needle. The vitelline membrane was peeled back and cuts were made at the interface between the spinal cord region of the neural tube and posterior somites and anterior presomitic mesoderm. Somites and presomitic mesoderm were peeled away to leave the posterior spinal cord unconfined on its sides. Eggs were then resealed and incubated overnight. Embryos that showed a good degree of healing and no obvious spinal cord ruptures were included in analysis.

### Grafting experiments

Small patches of dorsal hindbrain or spinal cord cells from GFP positive HH10-11 embryos were transplanted into the posterior spinal cord of stage-matched wild type embryos. A sharpened tungsten needle was used to peel back the vitelline membrane of donor and host embryos and to dissect out a small patch of dorsal cells from either the hindbrain or spinal cord. Tissue patches were transferred using a Gilson pipette and Ringers solution to the host embryos. A small incision was made into the posterior host embryo dorsal neural tube and tissue patches were pushed into this incision. Embryos were cultured *in ovo* overnight.

### Ferrofluid droplet experiments

Ferrofluid oil droplets were injected into the forebrain region of HH10-12 stage embryos using a Nanojet injector set to inject a volume of either 1, 2, or 4nl. Droplets were positioned in the lumen of either the hindbrain or spinal cord using a magnet and custom-built embryo holder. Embryos were fixed approximately 1.5 hours post droplet injection prior to immunostaining and confocal imaging or imaged live. Dorsal roof tissue thickness was measured at the point of maximum roof deformation in the hindbrain and spinal cord of injected embryos at the droplet location. Uninjected embryos at equivalent stages were measured at the same anterior-posterior level as controls.

### Atomic force microscopy

Atomic force microscopy (AFM) indentation measurements were performed using the setup described in ^44^. Briefly, HH10-12 stage chicken embryos were cultured on agarose plates and the vitelline membrane was removed to expose the embryo’s dorsal hindbrain and spinal cord. 89.3 µm diameter polystyrene beads were glued to Arrow-TL1 cantilevers (nominal spring constant = 0.01 N/m; NanoWorld, Neuchâtel, Switzerland) as probes. During measurements, the cantilever was approached with 10 µm/s until an indentation force of either 50nN or 75nN was reached. This force was maintained for 3 seconds, thus enabling the acquisition of the deformation over time (‘creep’).

In order to extract fluidity and elastic modulus values from the force-distance curves, ^45^ introduced the following piecewise model for the force ramp:

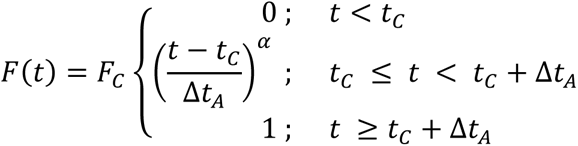

where *F_C_* is the magnitude of the applied force during the creep measurement reached in time *Δt_A_* and applied for time *t_C_*. α describes the shape of the force ramp (1 < *α* < 2), where α = 1 corresponds to a linear force ramp and α closer to 2 to a polynomial force ramp. Fitting this equation yields *F_C_*, *t_C_*, *Δt_A_*, and *α*. These values then derive the following equation for the indentation *δ*, here modified by substituting *E_0_*/(1-ν^2^) with the reduced instantaneous elastic stiffness, *k_0_*

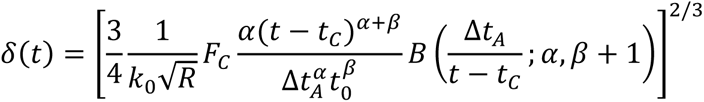

where *R* is the radius of the bead, *B()* is the incomplete beta function and the exponent, and β is the fluidity, which can have values ranging from 0 (for an elastic solid) to 1 (for a viscous fluid). Mean values for *k_0_* and β were determined from measurements taken across the hindbrain and along the spinal cord. Values obtained for *k_0_* were close to values obtained for the reduced apparent elastic modulus *K* = *E_0_*/(1-ν^2^) obtained by fitting the Hertz model 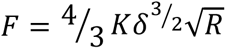 to the instantaneous indentation data (Figure. S2).

### MMP inhibitor treatment

After removal of the vitelline membrane covering the brain, 20µl of 1000µM broad-spectrum MMP inhibitor (GM6001) or DMSO diluted in Ringers solution, was added to the dorsal surface of HH10-11 stage embryos *in ovo.* Embryos were incubated overnight.

### Immunostaining

After fixing embryos overnight in 500µl 4% PFA at 4°C, they were washed 3X for ∼20 minutes per wash in 0.5% Triton-X-100 (Sigma-Aldrich, X100) in PBS (PBST). Embryos were incubated in blocking solution consisting of 4% Donkey serum in PBST for 1 hour at room temperature. Primary antibodies were diluted in blocking solution and incubated with embryos for 2 days at 4°C. After washing embryos in PBST ∼4 times over the course of 2 hours, secondary antibodies, Hoecst and Phalloidin were diluted in blocking solution and incubated with embryos for another 2 days at 4°C. Finally, embryos were washed in PBST prior to imaging.

### Primary and secondary antibodies used

Laminin (DSHB, 3H11) 1:100, Snail2 (Cell Signaling Technology, 9585S) 1:200, pMLC2 (Cell Signaling Technology, 3671S) 1:100, Hoecst 1:1000, Phalloidin 647 (Thermo Scientific, A22287) 1:500, Donkey Anti-Mouse 488n IgG (Abcam, ab150105), Donkey Anti-Rabbit 594 (Abcam, ab150076)

### Confocal imaging

Embryo cross sections and wholemounts were mounted on glass-bottom confocal imaging dishes in VECTASHIELD (Vector Labs, H-1000-10) mounting medium. Z-stacks were acquired at either 0.3 or 0.5µm z-steps using either an SP8 (Leica) or a SoRa Spinning Disk (Nikon) using either a 10X, 20X or 40X objective.

### Data analysis

All data analysis was performed in Python (www.python.org), with data stored in Excel. Boxplots were obtained using the seaborn library and show the median, upper and lower quartiles and whiskers extending to 1.5 times the interquartile range. Scatterplots were plotted using matplotlib. Statistical analysis was performed using the scipy.stats Python library. Paired t-tests were used to calculate p-values for the AFM data. Independent t-tests were used to calculate all other p-values. All tests were two-sided and data checked for normality using the Shapiro-Wilk test. In all cases P<0.05 was taken as the threshold for significance. Measurements were acquired from distinct samples in all cases unless otherwise stated. Details of measurements of reported features are listed below:

1. Tissue shape analysis. The thickness profile of neural tube cross-sections was acquired by measuring the radial tissue thickness along the lumen circumference starting from the approximate mid-point of the dorsal hindbrain roof. For wildtype characterisation, the distance along the lumen circumference was normalised for all samples so that tissue thickness at distance ‘0’ fell at the dorsal mid-point and tissue thickness at distance ‘1’ at the ventral midpoint of the neural tube cross section. A 4^th^ degree polynomial was fit to the thickness profiles to get an ‘average’ thickness profile for the hindbrain and spinal cord at HH11 and HH16. Images acquired by confocal microscopy were visualised using either ImageJ^46^ or napari^47^. Cross sections were either visualised directly, or by reordering the axes of image data acquired from wholemount samples in napari. For intubation experiment analysis the mean thickness of the first 100µm of dorsal roof tissue from the dorsal midpoint was compared. For BDX experiment analysis the mean thickness of the first 300µm of dorsal roof tissue from the dorsal midpoint was compared. Equivalent positions along the hindbrain were compared for assessing hindbrain roof thickness in MMP treated and control embryos.
2. Actin intensity along the apical surface. Curves were drawn along the apical surface of neural tube cross sections starting from the dorsal midpoint and moving ventrally. The actin intensity along the curve was measured using the ImageJ PlotProfile function. Normalised average actin intensities along the apical surface were obtained for each embryo and binned into three groups starting from the dorsal-most position and moving more ventrally. The mean actin intensity for each embryo within each bin was plotted.
3. Somite removal analysis. Z planes were selected in the region where somite removal had been performed and in a corresponding location in controls. Roof thickness was measured at the dorsal midpoint for z planes from each embryo.

### Data availability

Code used for data analysis will be made available on a GitHub repository and source data will be made available.

## SUPPLEMENTAL TEXT

### Supplementary theory note

In this Supplementary theory note, we propose a rheological model of neural tube expansion, including two possible deformation mechanisms: elastic stretching and viscous flowing, taking into account the geometrical differences between the hindbrain and the spinal cord. We also provide an approach to quantify the rheological parameters of tubular tissues by tracking the shape evolution of injected ferrofluid droplets. Based on the quantification of rheological parameters, we show that luminal fluid pressure can play a significant role in driving viscous long-term hindbrain expansion (Figure 1) and identify a ∼7-fold difference in long-term tissue viscosity between the hindbrain and the spinal cord. Finally, a more detailed model considering the dorsal-ventral inhomogeneity in tissue viscosity can well reproduce the dorsal thinning and elongation observed in experiments, further supporting the rheological mechanism we proposed.

#### 1. Rheological model of neural tube

The neural tube epithelium is treated as a viscoelastic Maxwell medium, which acts as an elastic solid on short time-scales and a viscous fluid on long time-scales ^48–51^. In the “spring-dashpot” representation (Figure ST1), the Maxwell medium behaves as a Newtonian fluid (viscous dashpot with viscosity *η*) in series with a Hookean solid (elastic spring with elastic modulus *E*) so that the total deformation is the sum of the elastic and viscous contributions.

**Figure ST1.**
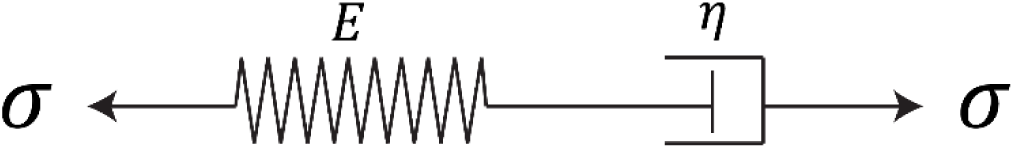
Schematic representation of the Maxwell rheological model.

Driven by luminal fluid pressure or other forces (such as the mechanical interaction with the injected ferrofluid droplet in the lumen, see Figure 2D for instance, the tissue deformation includes two parts: elastic deformation and viscous flowing. Several considerations argue in favour of considering a long-term viscous rheology. Firstly, the neural tube (including both the hindbrain and spinal cord) shows only moderate morphological changes on a short time-scale (seconds to minutes), with the expansion effects of ferrofluid droplets occurring on long time scales of hours (Figure S2F, see also section 2 below). Secondly, a number of experimental and theoretical works have shown that epithelial tissues are well-described by long-term viscosity, due to multiple active events resulting in local remodelling (e.g. divisions and intercalation)^48,49,52,53^. Thirdly, the luminal fluid pressure we measured is much lower than the expansion pressure engendered by the injected oil droplet, as well as orders of magnitude below epithelial stiffness, so that luminal fluid pressure is unlikely to lead to large elastic stretches during normal morphogenesis. In the modelling, we thus consider small elastic deformation characterized by elastic strain *∈*_*⍺*_, with the subscript *⍺* = *z*, *θ*, *r* respectively corresponding to the direction along the A-P axis, the azimuthal direction, and the radial direction.

Tissue deformation during neural tube morphogenesis can be characterized by comparing the current shape and the initial shape before the accumulation of luminal fluid (around Hamburg-Hamilton (HH) stage 11). The stretch ratio *λ*_*⍺*_ (*⍺* = *z*, *θ*, *r*), denoted as the ratio of the current length to the initial length of a tissue element along direction *⍺*, can evaluate the three-dimensional local deformation of the tissue. The stretch ratio *λ*_*⍺*_ of a viscoelastic tissue is contributed by both the elastic and viscous deformation. Consider the stretch ratio after elastic deformation is 1 + *∈*_*⍺*_, and denote the stretch ratio with only viscous flowing as *q*_*⍺*_. Then the total stretch ratio *λ*_*⍺*_ is the product of these two parts:

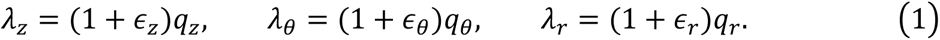

The small elastic deformation of the tissue follows the linear elastic strain-stress relation, i.e. Hooke’s law (Landau et al., 1986):

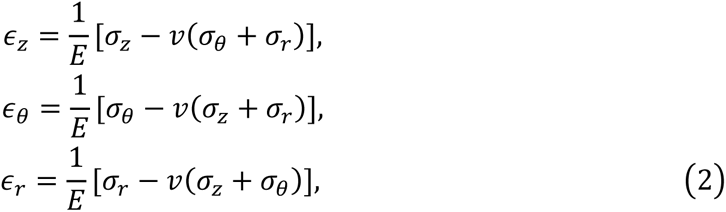

with *E* the Young’s modulus of the tissue, *v* the Poisson’s ratio, and *σ*_*⍺*_ the elastic stress in direction *⍺*. Given that volume changes in cells involve forces much larger than those at play for morphogenetic shape changes^54^, soft tissues are usually modelled as incompressible elastic media, which corresponds to *v* = 1/2.

As for Newtonian fluids, the rate of viscous deformation is proportional to shear stress. This relation can be extended to three-dimensional and large viscous deformation (see ^55^ for the theoretical framework of large viscoelastic deformation):

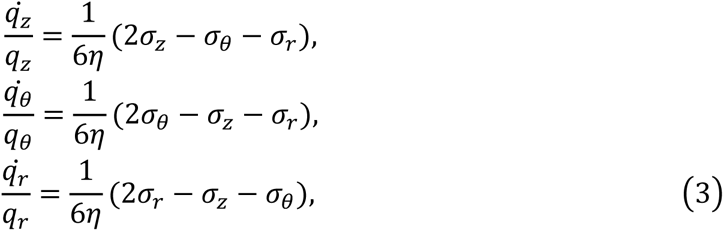

with *η* the tissue viscosity. The stress *σ*_*⍺*_ in Eqs. (2) and (3) can be obtained by discussing the force balance condition in direction *⍺*.

#### 2. Quantification of rheological parameters

Although experimentally measuring long-term tissue mechanical properties has been an historic challenge in the field, the shape change of ferrofluid droplets inside the tubular tissues provides a way to clarify the deformation mechanism and infer tissue rheology parameters (such as viscosity *η* and elastic modulus *E*)^50,56^. In this part, we provide theoretical foundations for the quantification of mechanics in tubular tissues.

##### 2.1. Shape – pressure relationship of ferrofluid droplet

After being injected into the lumen of the neural tube, a ferrofluid droplet will deform into an ellipsoid due to the confinement from the tissue wall (Figure ST2). Its surface tension *γ* is balanced with the inner pressure *p*_*d*_ and an outer pressure, which is the luminal fluid pressure *p*_*l*_ in its polar region and tissue-droplet interfacial pressure *p* in its equator. The force balance in the fluid droplet also depends on the local curvatures:

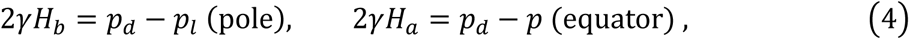

with *H*_*b*_ and *H*_*a*_ respectively the mean curvatures in the pole and the equator (Figure ST2).

The above force balance links the tissue-droplet interfacial pressure *p* and the droplet inner pressure *p*_*d*_ with the droplet shape (surface curvature):

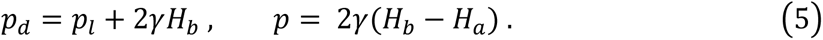

For an ellipsoidal droplet with *a* and *b* respectively the lengths of its short and long axes, we have *H*_*b*_ = 2*b*/*a*^2^ and *H*_*a*_ = *a*/*b*^2^ + 1/*a* ^56^. Using Eq. (5), the pressures *p* and *p*_*d*_ can be inferred from the lengths of two major axes (i.e. *a* and *b*) of the ferrofluid droplet.

**Figure ST2.**
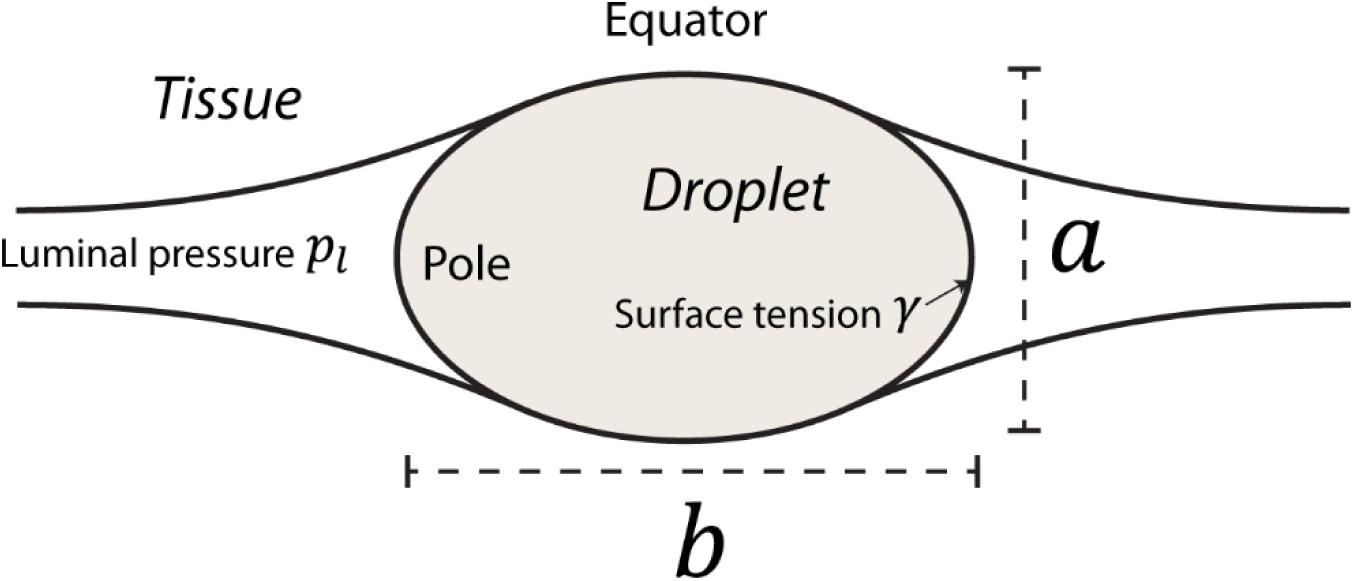
Schematic geometry and mechanics of a ferrofluid droplet injected into the tissue lumen. The droplet with surface tension *γ* is deformed as an ellipsoid with short-axis length *a* and long-axis length *b*. Its equator (and nearby region) is in contact with the tubular tissue, while its pole is pressurized by luminal fluids (*p*_*l*_).

##### 2.2. Tissue viscosity of hindbrain inferred from droplet shape evolution

The hindbrain can be considered a thin-walled tissue, whose tissue thickness is smaller than tissue radius (approximately two or three times smaller than the tissue radius in HH11 and becomes approximately five times smaller than the radius in HH16, see Figure 1D for snapshots of the tissue cross-sections). This allows us to approximate the hindbrain tissue as a viscoelastic membrane, where the stress in the radial direction *σ*_*r*_ is much smaller than the in-plane stresses (*σ*_*θ*_ and *σ*_*z*_) and can be neglected. Thus, the deformation of a thin-wall tissue like the hindbrain is determined by the in-plane forces. Using the fact that the in-plane stresses in a thin wall tissue are homogeneously distributed along the thickness/radial direction, we can define in-plane tissue tensions *T*_*θ*_ = *hσ*_*θ*_ and *T*_*z*_ = *hσ*_*z*_ (*h* is the tissue thickness), which are the sum of in-plane stresses (*σ*_*θ*_ and *σ*_*z*_) along the tissue thickness, and share the same unit (force per unit length) with the surface tension of ferrofluid droplet *γ* ^57–60^. In this way, the interaction between the ferrofluid droplet and the viscoelastic hindbrain can be modelled as a two-layer membrane system (Figure ST3): a viscoelastic membrane with in-plane tension *T*_*θ*_ and *T*_*z*_, and a liquid membrane with surface tension *γ*.

**Figure ST3.**
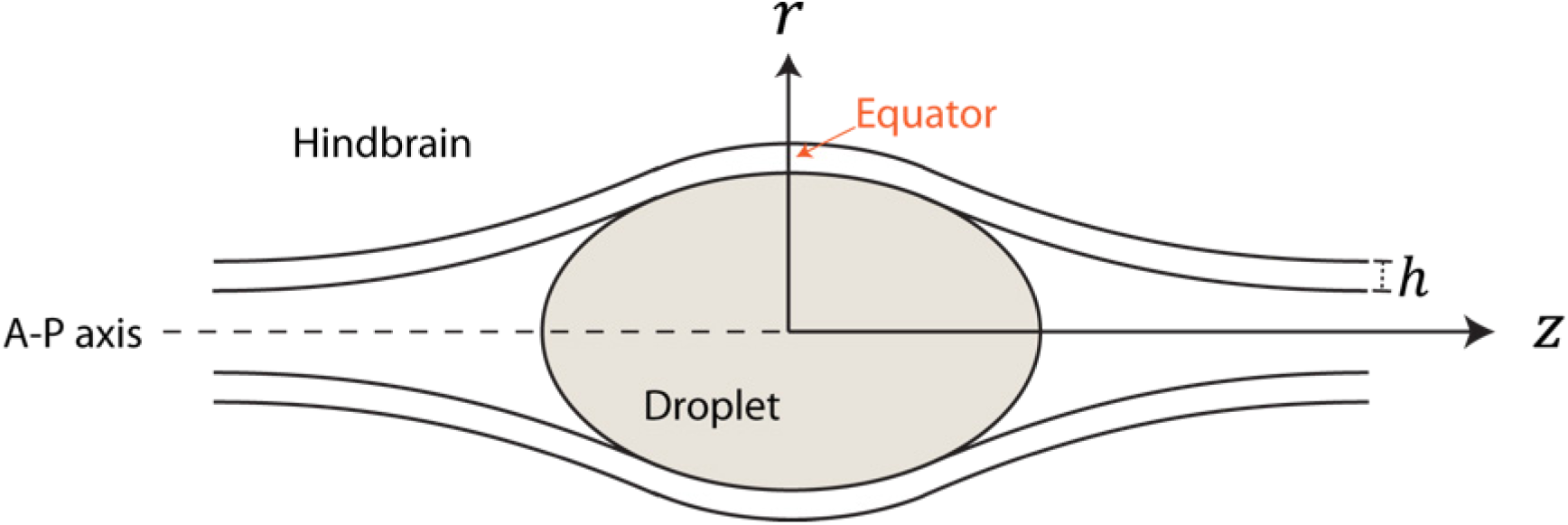
Schematic of the hindbrain-droplet system.

The hindbrain-droplet system is cylindrically symmetric to the A-P axis. Note that in this geometry (Figure ST3), the axial stretching force *T*_*z*_(*σ*_*z*_) and corresponding deformation (e.g. *λ*_*z*_ and *q*_*z*_) is in the meridional direction, not necessarily parallel to the A-P axis (i.e. z-axis). In such configuration, we have the azimuthal stretch ratio *λ*_*θ*_ = *r*/*R*, with *r* and *R* respectively the current and initial radii of the hindbrain^61,62^. Importantly, the middle of the contact region (i.e. the droplet equator, see Figure ST3) of the hindbrain-droplet system satisfies a simple geometric relation: the current tissue radius is half length of the droplet’s short axis (that is *r* = *a*/2), which directly links the tissue deformation and droplet shape.

Importantly, the shape evolution of the ferrofluid droplet depends on tissue viscosity. This is made clear from the fact that we observe droplet rounding to occur over time scales of hours, many orders of magnitude longer than what would be expected from rounding in water due to surface tension (Figure S2F). Thus, we can use the dynamics of droplet rounding as a proxy to estimate the long-term viscosity of the tissue, at the time scales of hours which are relevant for morphogenesis.

Considering the geometric relation *λ̇*_*θ*_/*λ_*θ*_* = *ȧ*/*a*, and small elastic strains *∈*_*⍺*_ in Eq. (1), which leads to *λ̇*_*θ*_/*λ*_*θ*_ ≈ *q̇*_*θ*_ /*q*_*θ*_, we can rewrite the second formula in the evolution equation (3) as

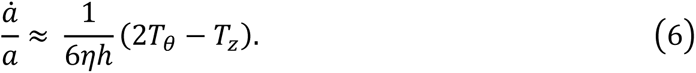

Eq. (6) clearly shows the dependence of droplet shape on tissue viscosity *η* and in-plane tissue tensions *T*_*θ*_ and *T*_*z*_, which can be obtained by discussing the force balance in the equator position (Figure ST3).

Similar to the force balance between surface tension and pressure in liquid droplets (e.g. Eq. (4)), the in-plane tissue tensions of the hindbrain are balanced with the interfacial pressure *p*. Besides, in the meridional direction (i.e. z-direction in the droplet equator, see Figure ST3), the in-plane tissue tension *T*_*z*_ and droplet surface tension *γ* are balanced with the inner pressure of the droplet *p*_*d*_. These lead to the following relations in the droplet equator:

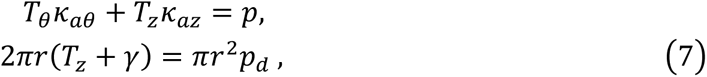

with the principal curvature in the azimuthal direction ***κ***_*aθ*_ = 2/*a*, in the meridional direction ***κ***_*az*_ = 2*a*/*b*^2^, and tissue radius *r* = *a*/2 as aforementioned.

We introduce the aspect ratio of ferrofluid droplet *s* = *b*/*a*, so that the combination of Eqs. (5–7) gives the shape evolution equation of the ferrofluid droplet:

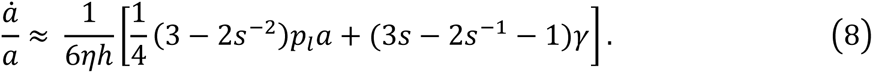

Considering the luminal fluid pressure *p*_*l*_∼15 Pa (Figure 1C), lumen diameter in the hindbrain (or short-axis length of ferrofluid droplet) *a*∼200 μm, and the surface tension of ferrofluid *γ*∼0.026N/m, one can find *p*_*l*_*a*/4 << *γ*, and thus that the first term in the right side of shape evolution equation (8) can be neglected.

Note that the axis lengths *a* and *b* are not independent, but obey the geometric constraint of constant droplet volume *V*_*d*_ = (4*π*/3)*a*^2^*b*. With this constraint, the droplet shape (evolution) can be evaluated by a single shape parameter (*a*, *b*, or aspect ratio *s*), and these parameters are related to each other: for instance, we have

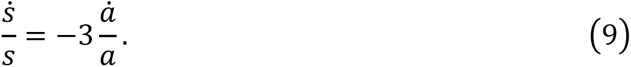

By Eq. (9), we can replace the rate of shape evolution *ȧ*/*a* with *ṡ*/*s*, and rewrite the shape evolution equation (8) as

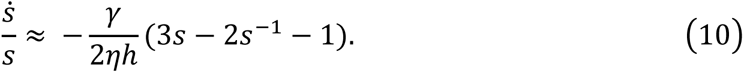

In Eq. (10), the tissue thickness also evolves with time, satisfying *ḣ*/*h* = *λ̇*_*r*_/*λ*_*r*_ ≈ *q̇*_*r*_ /*q*_*r*_. Combined with the third formula in evolution equation (3), we can get the evolution law of tissue thickness. However, thickness evolution is quite slow compared with the shape change of the ferrofluid droplet, thus the tissue thickness *h* in Eq. (10) can be treated as a constant (which would bring quite small errors as shown in Figure ST5(a) but allows us to get analytic solutions). Then we can easily get the analytic solution of Eq. (10) as

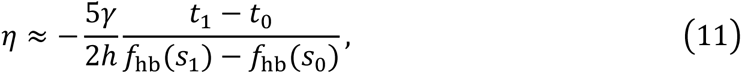

with 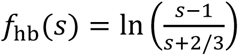 *s*_0_ and *s*_1_ respectively the aspect ratios of ferrofluid droplet in time points *t*_0_ and *t*_1_.

##### 2.3. Tissue viscosity of spinal cord inferred from droplet shape evolution

Unlike the hindbrain, the spinal cord has a tissue thickness comparable with or even larger than its lumen radius. Besides, Figures 2B-C showed that the removal of surrounding tissues can enlarge the spinal cord, indicating the spinal cord is mechanically confined by surrounding tissues (such as somites). Thus, to discuss how the ferrofluid droplet mechanically interacts with the spinal cord, it is more realistic to consider its interaction with a broader mechanical surrounding (including the spinal cord and the surrounding tissue, see Figure ST4), compared to the thin-walled approximation employed to the hindbrain (see Subsection 2.2 for details). Let *r*_*i*_ and *r*_*o*_ respectively denote the inner and outer radius of the tissue system, we have *r*_*o*_ ≫ *r*_*i*_ = *a*/2. In this scenario, the stress in A-P direction *σ*_*z*_, which arises from droplet pressure *p*_*d*_ or luminal fluid pressure *p*_*l*_ is quite small, thus the tissue deformation is mainly driven by the azimuthal stress *σ*_*θ*_ and the radial stress *σ*_*r*_.

**Figure ST4.**
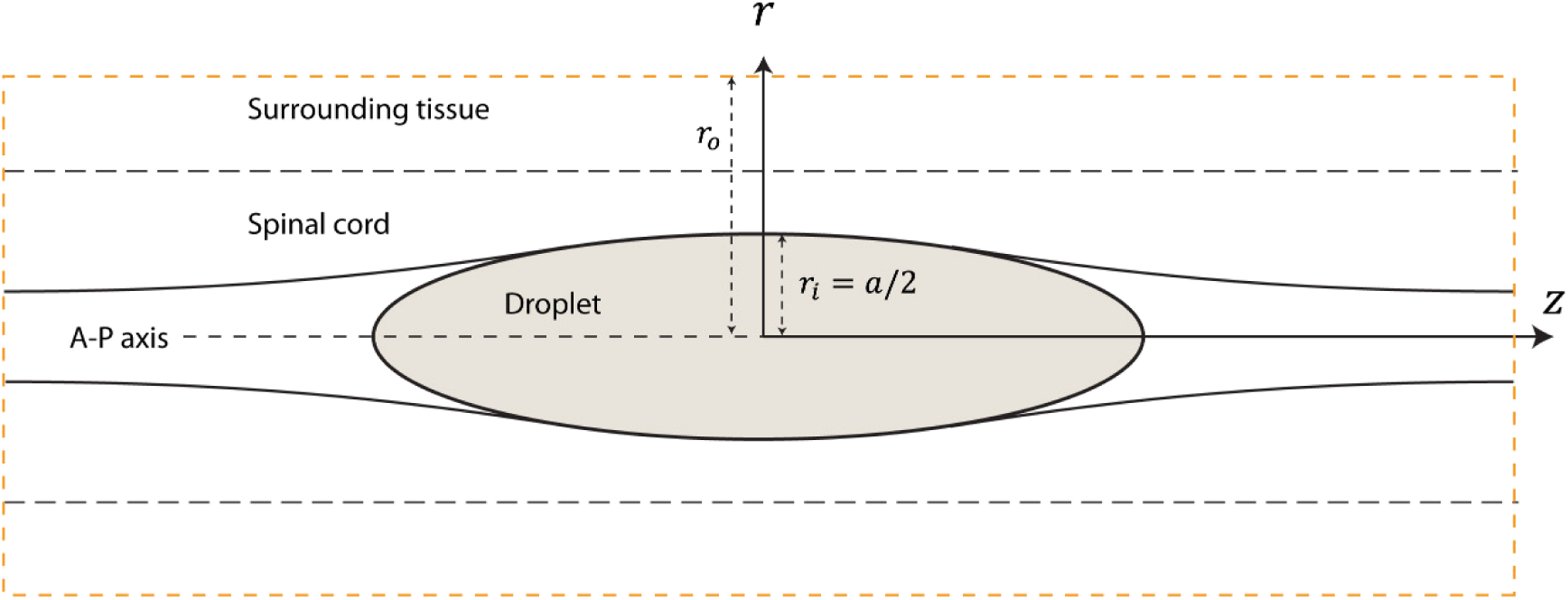
Schematic of the spinal cord-droplet system.

Similar to the discussion in Subsection 2.2, we can get *λ̇*_*θ*_/*λ*_*θ*_ = *ȧ*/*a* at *r* = *r*_*i*_ = *a*/2, and *λ̇*_*θ*_/*λ*_*θ*_ ≈ *q̇*_*θ*_ /*q*_*θ*_ for small elastic strains. Using the evolution law of azimuthal viscous deformation in Eq. (3), we have

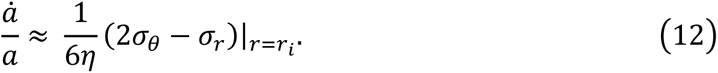

The tissue stresses *σ*_*θ*_ and *σ*_*r*_ are generated by the interfacial pressure *p* and radially distributed as ^63^

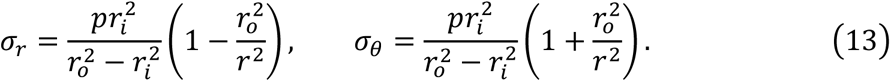

Considering *r*_*o*_ ≫ *r*_*i*_ leads to (2*σ*_*θ*_ − *σ*_*r*_)|_*r*=*r*_*_*i*_* ≈ 3*p* in Eq. (11). The interfacial pressure *p* can be inferred from the droplet shape by relation (5). Then, we can rewrite Eq. (12) as

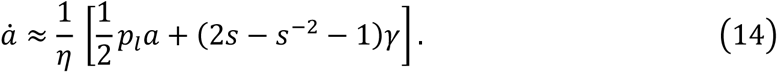

The first term in the right of Eq. (14) can be neglected in data fitting, based on the scaling analysis that *p*_*l*_*a* is an order of magnitude smaller than droplet surface tension *γ*. Although the combination of Eqs. (9) and (14) can give close solutions of the shape evolution of ferrofluid droplet, they have to be solved numerically. Submitting Eq. (9) into Eq. (14), the shape evolution equation becomes:

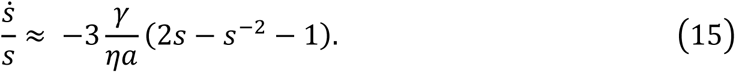

Eq. (9) indicates the aspect ratio *s* evolves faster than the short-axis length *a*, which allows us to approximate *a* as a constant. In this way, we can get the analytic formula for the data fitting of tissue viscosity of Eq. (15), which could be used to directly infer the tissue viscosity of the spinal cord from the shape evolution of droplet:

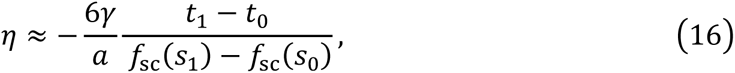

with 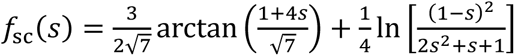. The complete solution of Eqs. (9) and (14) indicates the analytic formula (16) is accurate enough (Figure ST5(b)).

**Figure ST5.**
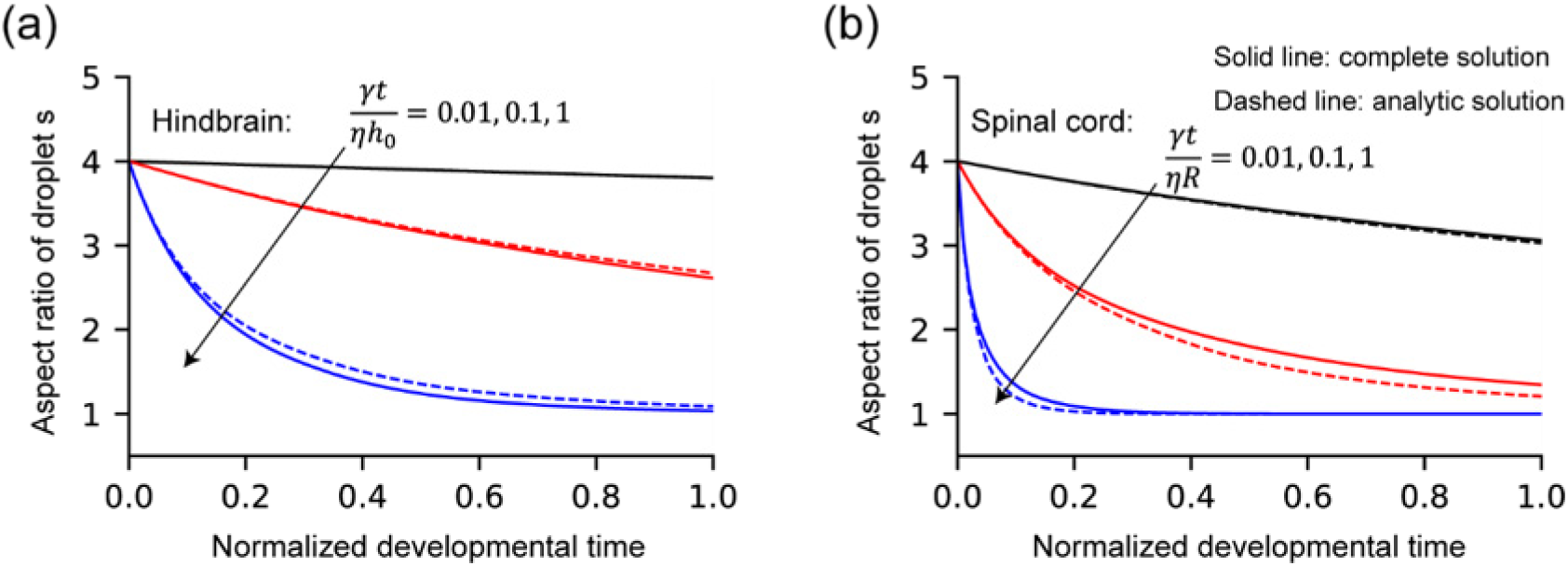
Comparison between the complete and approximated solutions of the droplet shape evolution. (a) Time evolution of the aspect ratio of ferrofluid droplet in the hindbrain. Analytic solution in Eq. (11) treating tissue thickness *h* as a constant (i.e. *h* = *h*_0_) is compared with the complete solution with thickness *h* also evolving. (b) Time evolution of the aspect ratio of ferrofluid droplet in the spinal cord. Analytic solution in Eq. (16) considering lumen diameter (or short-axis length of droplet) *a* as a constant is compared with the complete solution.

#### 3. Tissue deformation in normal development

Experimentally, we found that luminal fluids play an important role in the development of the neural tube, and the variation in luminal fluid pressure greatly affects the hindbrain deformation and morphogenesis (characterized by a thinning of the dorsal tissue and expansion of the lumen, see Figure 1). Here, we quantitatively discuss if the morphogenesis of the hindbrain could be a result of viscous or elastic deformation driven by luminal fluid pressure.

Importantly, we observe droplet rounding over hours after injection into the lumen of neural tube (Figure S2F), which is many orders of magnitude longer than what would be expected for droplets alone under surface tension (i.e. free from tissue confinement). This hints at a long-term viscous relaxation of tissue on the timescale of hours. Quantitatively, we tracked the tissue deformation rate *ȧ*/*a* (with *a* the tissue diameter and *ȧ* its rate) after injecting the ferrofluid droplet.

Driven by the droplet pressure, the hindbrain diameter increases by approximately 1.5X in 1-2 hours, suggesting the deformation rate *ȧ*/*a*∼h^−1^. The viscous deformation rate of the hindbrain is proportional to in-plane tissue tension (Eq. (6)). The in-plane tension of the hindbrain expanded by the droplet is in the same order with the droplet surface tension *γ*, while in the normal development, the hindbrain is expanded by the luminal fluid pressure *p*_*l*_ and the tissue tension is ∼*p*_*l*_*a* based on Young-Laplace law (Fig. 2A) and Eq. (8). With the droplet surface tension *γ*∼0.026N/m, luminal fluid pressure *p*_*l*_∼15Pa and hindbrain diameter *a*∼200μm, we found *p*_*l*_*a* is an order of magnitude smaller than *γ*, thus would speculate the viscous deformation rate in the normal development is an order of magnitude slower than the scenario with ferrofluid droplet, that is *ȧ*/*a*∼0.1h^−1^, which would leads to several folds of hindbrain expansion in around 20 hours (from HH11 to HH16), consistent with experimental observations (Figure 1).

Tissue viscosity *η*, the rheological parameter characterizing the tissue resistance to viscous flowing, can be inferred from the droplet shape evolution, as discussed in Subsection 2.2 and 2.3. Analytic formulas (11) and (16) provide a simple way to estimate the tissue viscosity *η* by comparing the aspect ratios in two different timepoints (usually with a time interval of one or two hours in our experimental setting). We estimate the tissue viscosity in the hindbrain as 2.3±0.84 kPa · h (mean ±SD), and that in the spinal cord as 17 ±10 kPa · h. Importantly, our quantification shows that, the hindbrain is more fluidized and easier to deform than the spinal cord under mechanical loading. This is in consistent with atomic force microscopy (AFM) measurement (Figure 2H) and experimental evidence supporting tissue fluidization in the hindbrain (Figure 3 and 4).

The minimal deformation of the hindbrain tissue observed immediately following droplet placement suggests a small degree of elastic deformation. In the normal development with the hindbrain expansion driven by luminal fluid pressure, the elastic deformation of the tissue would be even smaller, far less than enough to reproduce the several folds of hindbrain expansion observed in normal morphogenesis (Figure 1).

The viscous evolution laws (e.g. Eqs. (8) and (14)) in section 2 include both the contributions from the luminal fluids and ferrofluid droplet, while in the normal development, the luminal fluid pressure *p*_*l*_ is the only driving force for tissue deformation. After removing the impact of the droplet (we could set *γ* = 0 and *s* → ∞), the sole impact of the luminal pressure *p*_*l*_ on the viscous evolution or elastic displacement can be obtained:

- time evolution rate of tissue diameter *ȧ* due to tissue viscous deformation:

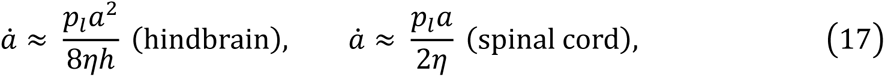

##### 4. Rheological inhomogeneity in the hindbrain

In the previous analysis and discussion, the tissue is considered to be axially varying (i.e. the geometric and mechanical properties may vary along A-P axis) but are circumferentially homogeneous. However, in the normal development, the dorsal and ventral regions of the hindbrain show distinct morphological evolution (Figure 1D). In the main text, this morphological difference along D-V axis is attributed to higher deformability of the dorsal tissue compared with its ventral partner (Figure 3 and 4). Tissue fluidization driven by neural crest ECM remodelling in the dorsal hindbrain would decrease the tissue viscosity and make the dorsal hindbrain easier to deform under mechanical load. To quantitatively test this rheological assumption, here we extend the model to consider the D-V inhomogeneity in tissue viscosity.

**Figure ST6.**
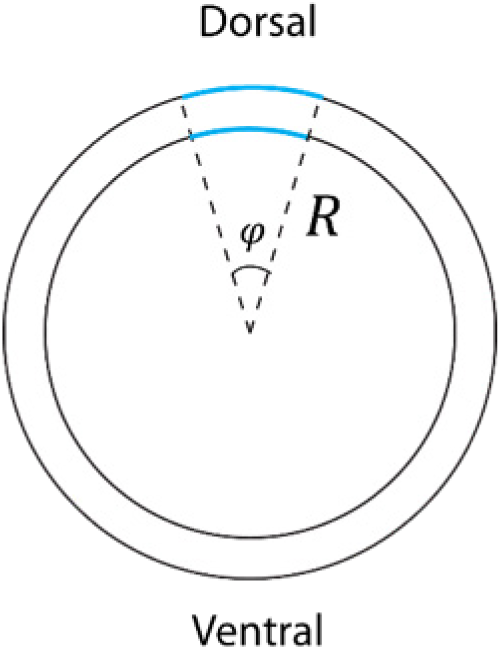
Schematic of the cross-section of the hindbrain.

In the cross-section of the hindbrain (see Figure ST6 for schematic), consider the dorsal and ventral regions respectively with initial arc lengths *L*_*D*_ = 2*πφR* and *L*_*V*_ = 2*π*(1 − *φ*)*R* (*φ* is the initial length ratio of the dorsal to ventral region), and the same initial tissue thickness *h*_0_, but different tissue viscosities, i.e. *η*_*D*_ < *η*_*V*_. As a viscoelastic membrane with deformation dominated by viscous flowing (see Subsection 2.2 for details), the time evolution of the stretch ratios of the hindbrain tissue yields

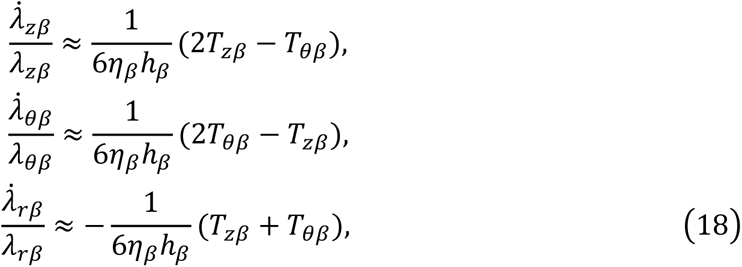

with *β* = *D*, *V* respectively standing for the dorsal and ventral region of the hindbrain.

The azimuthal stretch *λ*_*θβ*_ is the ratio of current arc length *l*_*β*_ to its initial value *L*_*β*_, that is *λ*_*θβ*_ = *l*_*β*_/*L*_*β*_. The regional arc lengths are related to the current tissue radius *r*, and should satisfy the geometric constraint: *l*_*D*_ + *l*_*V*_ = 2*πr*. We can easily get *λ̇*_*θβ*_/*λ*_*θβ*_ = *l*_*β*_/*l*_*β*_ in Eq. (23). Similarly, the radial stretch ratio *λ*_*rβ*_ evaluates the change of tissue thickness *h*_*β*_ and we have *λ̇*_*rβ*_/*λ*_*rβ*_ = *ḣ*_*β*_/*h*_*β*_. In the A-P axial direction, the deformations of the dorsal and ventral regions are synchronous: *λ*_*zD*_ = *λ*_*zV*_ and *λ̇*_*zD*_ = *λ̇*_*zV*_.

The in-plane tissue tensions are balanced with the luminal fluid pressure *p*_*l*_. The dorsal and ventral hindbrains sustain the same azimuthal tissue tension: *T*_*θD*_ = *T*_*θV*_ = *T*_*θ*_ = *p*_*l*_*r*, and their total axial forces (along A-P axis) are balanced with *p*_*l*_ · *πr*^2^: *T*_*zD*_*l*_*D*_ + *T*_*zV*_*l*_*V*_ = *p*_*l*_ · *πr*^2^.

It is convenient to non-dimensionalize the parameters: we have tissue geometric parameters *r̅* = *r*/*R*, *l̅*_*β*_ = *l*_*β*_/*R*, and *h̅*_*β*_ = *h*_*β*_/*h*_0_, normalized tissues tensions *T̅*_*θ*_ = *T*_*θ*_/(*p*_*l*_*R*), *T̅*_*zβ*_ = *T*_*zβ*_/(*p*_*l*_*R*). The evolution of the hindbrain morphology depends on two parameters: the viscosity inhomogeneity Λ = *η*_*D*_/*η*_*V*_ and a dimensionless time *r* = *pRt*_∗_/(6*η*_*V*_*h*_0_), with *t*_∗_ the total developmental time. Considering luminal fluid pressure *p*_*l*_ ≈ 15Pa, initial radius to thickness ratio *R*/*h*_0_ ≈ 2, the developmental time *t*_∗_ ≈ 20h and ventral tissue viscosity *η*_*V*_∼10^3^Pa · h, we can estimate *r*∼0.1. The final governing equation system includes three parts (with the uppers bars for quantities dropped):

- time evolution laws for arc lengths and tissue thicknesses:

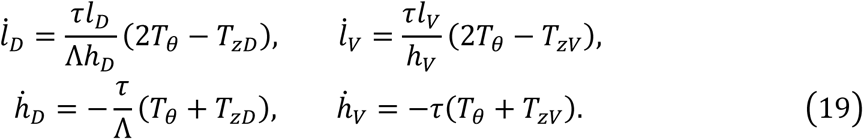

- force balance conditions:

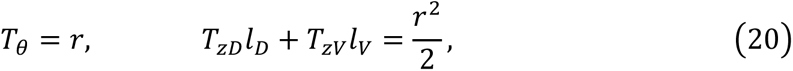

- geometric constraints:

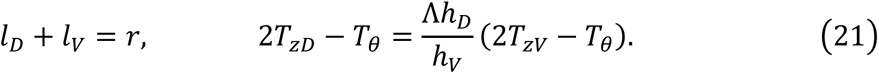

Interestingly, the rheological model considering a lower tissue viscosity in the dorsal hindbrain can well reproduce the morphological features observed in the morphogenesis of neural tube: the dorsal hindbrain becomes much thinner than the ventral hindbrain with development (Figure ST7(a)), meanwhile the arc length of the dorsal hindbrain elongates with time (Figure ST7(b)). This further validates that the rheological model can well describe the morphogenesis of neural tube and tissue rheological behaviours respond to the morphological features.

**Figure ST7.**
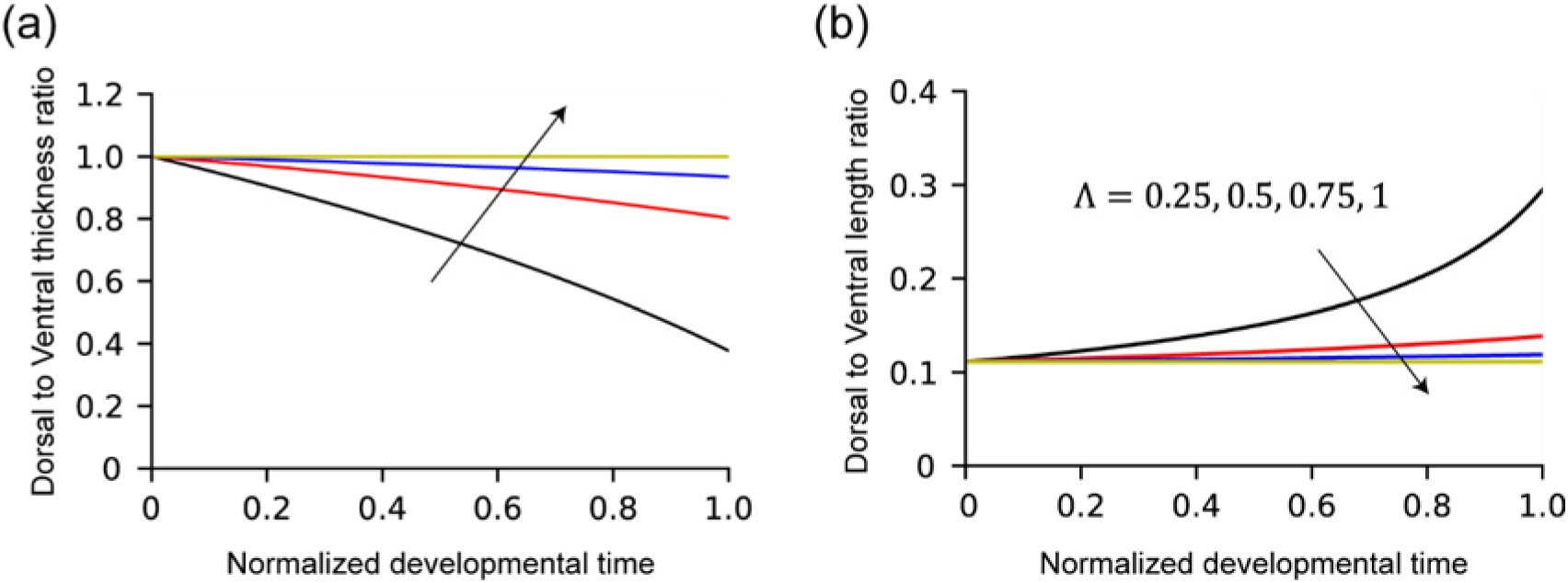
The time evolution of dorsal to ventral thickness ratio and length ratio with development. This numerical example sets the dimensionless parameter *r* = 0.1, and varies the viscosity inhomogeneity Λ.

---

**Figure S1.**
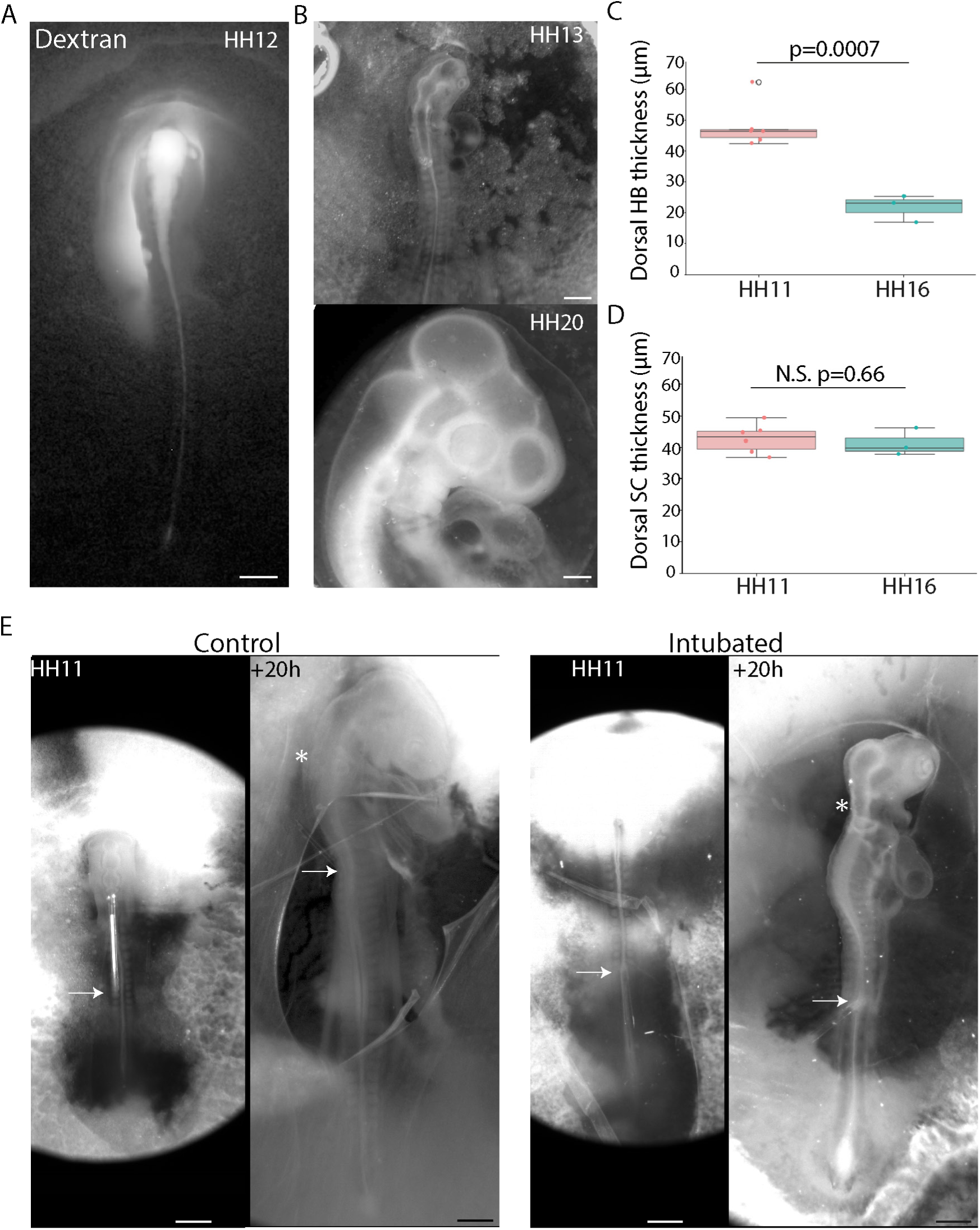
Neural tube characterisation and intubation experiments. (A) An HH12 stage embryo injected in the neural tube lumen with fluorescent Dextran. (B) Brightfield images of the anterior of embryos used in pressure measurements. (C) Mean thickness of the dorsal hindbrain in HH11 (n=6) and HH16 (n=3) stage embryos. (D) Mean thickness of the dorsal hindbrain in HH11 (n=6) and HH16 (n=3) stage embryos. (E) Brightfield images of control and intubated embryos just after intubation and 20 hours later. White arrows mark site of intubation. White asterisks mark the hindbrain. All scale bars are 500µm.

**Figure S2.**
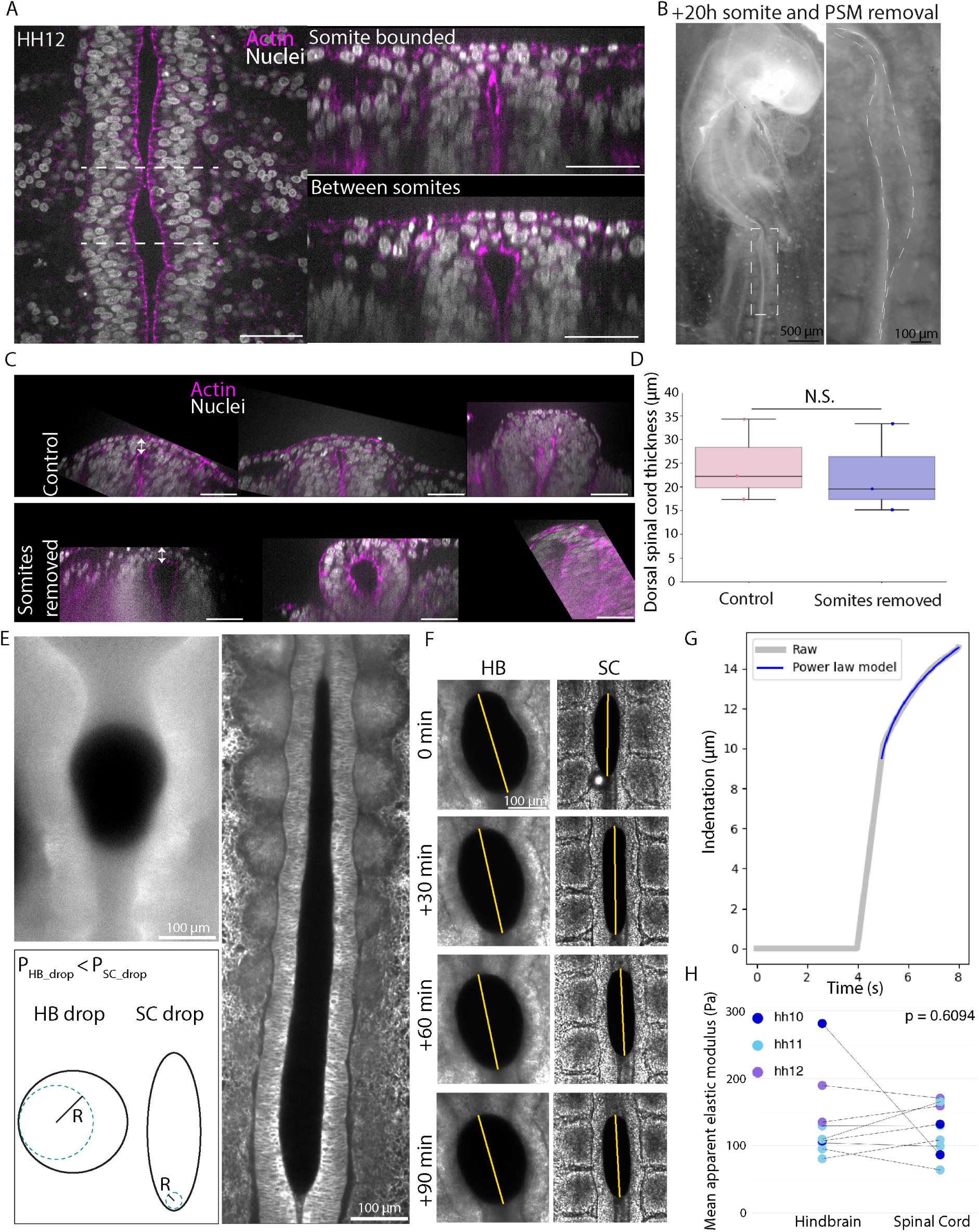
Geometry and tissue mechanics experiments. (A) Confocal images of dorsal and cross-sectional views of the spinal cord region in an HH12 stage embryo. Dashed lines indicate level of cross-sectional views. (B) A widefield image of an embryo ∼20 hours after posterior somite and PSM removal with zoomed-in view of the widened spinal cord region. (C) Confocal images of cross-sectional views of the spinal cord in embryos 20 hours after control cuts or somite removal. (D) Dorsal spinal cord thickness in control (n=3) and somite removed (n=3) embryos, measured at the dorsal midpoint. (E) Close up views of ferrofluid droplets in the hindbrain and spinal cord and a schematic illustrating the higher curvature at the droplet-lumen interface and thus higher pressure exerted by droplets in the spinal cord region compared to the hindbrain region. (F) Ferrofluid droplet rounding dynamics in the hindbrain and spinal cord. (G) Fitting of AFM indentation data to extract fluidity ‘β’. (H) Mean elastic modulus in the hindbrain and spinal cord of HH10-12 stage embryos (n=9). White scale bars are 50µm unless otherwise stated.

**Figure S3.**
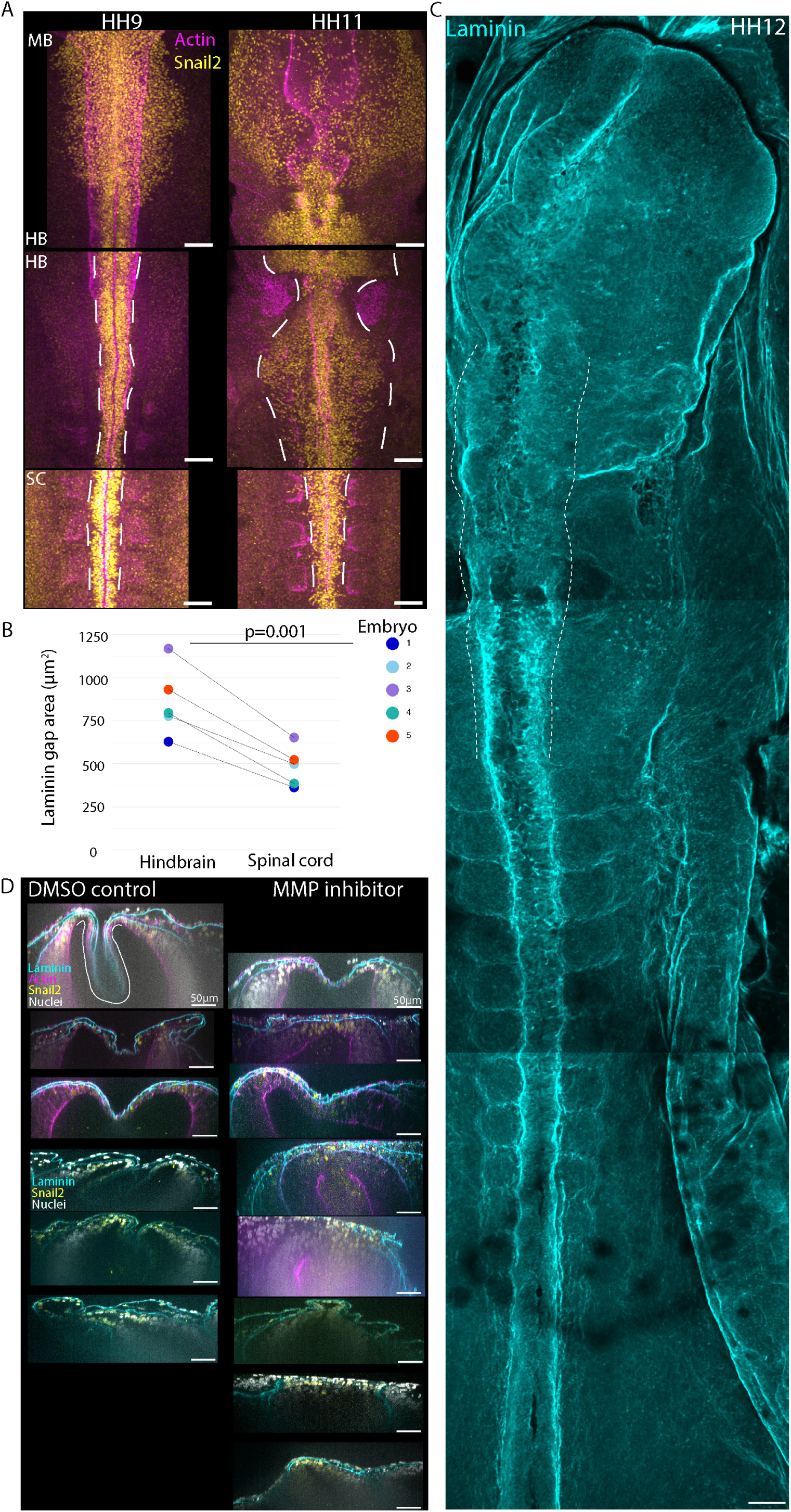
Neural crest and ECM experiments. (A) Confocal images showing Snail2+ cells at the dorsal surface of HH9 and HH11 stage brain and spinal cord regions. Dashed lines depict the extent of Snail2+ cell spreading away from the dorsal midline. (B) Total area of gaps in ECM on the dorsal surface of the hindbrain and spinal cord of pre-brain expansion embryos (n=5). (C) Confocal images stitched together to show a 3D rendering of the laminin matrix in a HH12 stage embryo. Dashed lines depict the boundaries of the hindbrain region. (D) Confocal images of DMSO and MMP inhibitor treated embryo hindbrains, ∼20 hours post treatment. Actin not stained for in bottom 3 images in control and inhibitor panels. White scale bars are 100µm unless otherwise stated.

**Figure S4.**
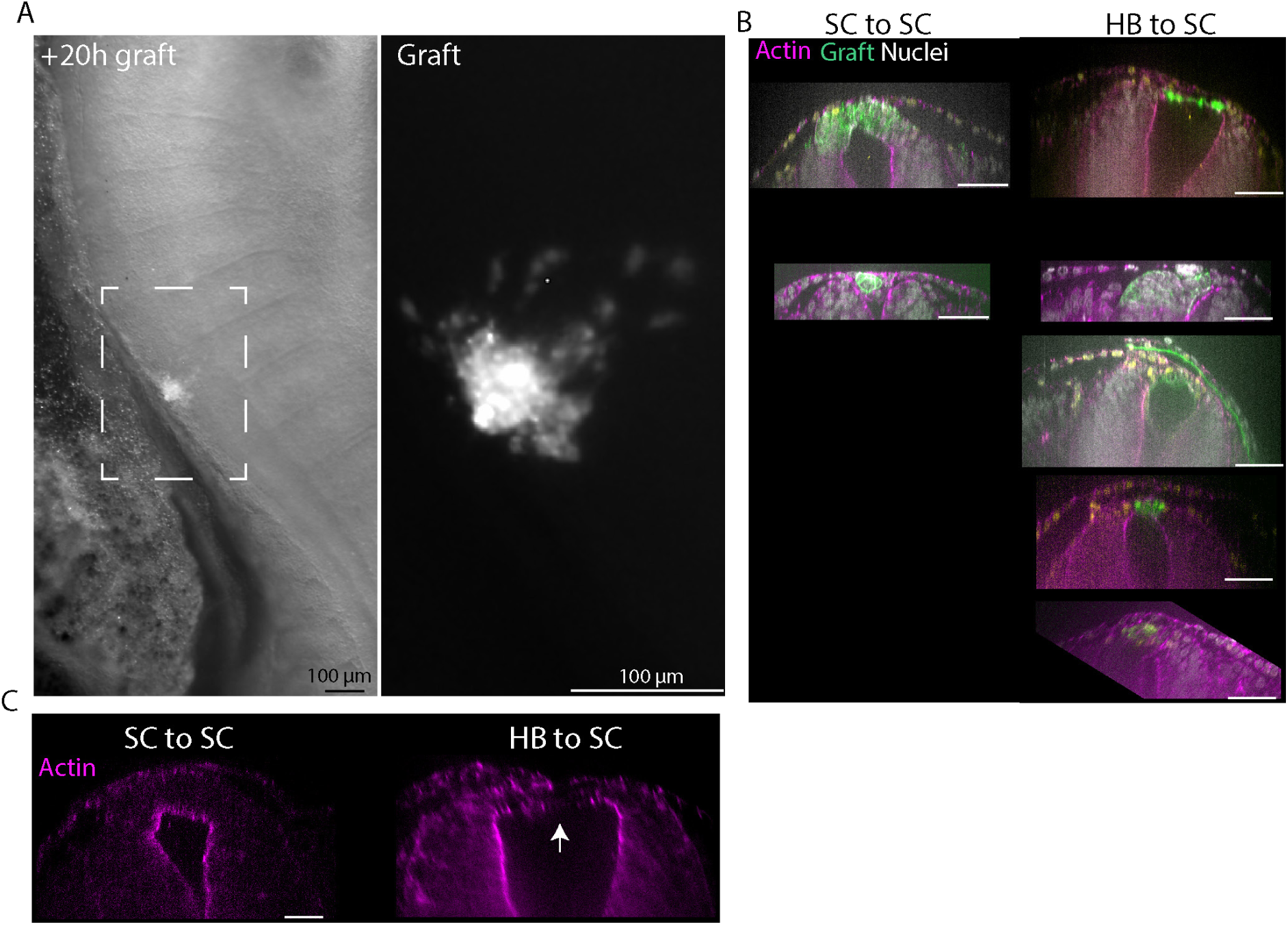
Graft experiments. (A) Widefield images of an embryo ∼20 hours after introduction of a GFP+ graft into the spinal cord and a zoomed-in view of the graft region. (B) Confocal images of cross-sectional views of the spinal cord at spinal cord-spinal cord and hindbrain-spinal cord graft sites. (C) Confocal images showing actin organisation at the apical lumen surface in a spinal cord-spinal cord graft region and a hindbrain-spinal cord graft region. White arrow indicates loss of apical actin localisation. White scale bars are all 50µm unless otherwise stated.

## Notes

### Competing Interest Statement

The authors have declared no competing interest.

### Summary of Updates

This version includes modelling and experimental approximation of viscous tissue deformation of the hindbrain and spinal cord under lumen pressure.

